# Live microscopy of multicellular spheroids with the multi-modal near-infrared nanoparticles reveals differences in oxygenation gradients

**DOI:** 10.1101/2023.12.11.571110

**Authors:** Angela C. Debruyne, Irina A. Okkelman, Nina Heymans, Cláudio Pinheiro, An Hendrix, Max Nobis, Sergey M. Borisov, Ruslan I. Dmitriev

## Abstract

Assessment of hypoxia, nutrients, metabolite gradients, and other hallmarks of the tumor microenvironment within 3D multicellular spheroid and organoid models represents a challenging analytical task. Here, we report red/near-infrared emitting cell staining O_2_-sensitive nanoparticles, which enable measurements of spheroid oxygenation on a conventional fluorescence microscope. Nanosensor probes, termed ’MMIR’ (multi-modal infrared), incorporate a near-infrared O_2_-sensitive metalloporphyrin (PtTPTBPF) and a deep red aza-BODIPY reference dyes within a biocompatible polymer shell, allowing oxygen gradients quantification *via* fluorescence ratio and phosphorescence lifetime readouts. We optimized staining techniques and evaluated nanosensor probe characteristics and cytotoxicity. Subsequently, we applied nanosensors to the live spheroid models based on HCT116, DPSCs, and SKOV3 cells, at rest and treated with drugs affecting cell respiration. We found that the growth medium viscosity, spheroids size, and formation method influenced spheroid oxygenation.

Unexpectedly, some spheroids (produced from HCT116 and dental pulp stem cells) exhibited ’inverted’ oxygenation gradients, with higher core oxygen levels than the periphery. This contrasted with the frequently encountered ‘normal’ gradient of hypoxia towards the core caused by diffusion. Further microscopy analysis of spheroids with an “inverted” gradient demonstrated metabolic stratification of cells within spheroids: thus, autofluorescence FLIM of NAD(P)H revealed the formation of glycolytic core, and localization of OxPhos-active cells at the periphery. Collectively, we demonstrate a strong potential of NIR-emitting ratiometric nanosensors for advanced microscopy studies targeting live and quantitative real-time monitoring of cell metabolism and hypoxia in complex 3D tissue models.

**Graphical abstract:** 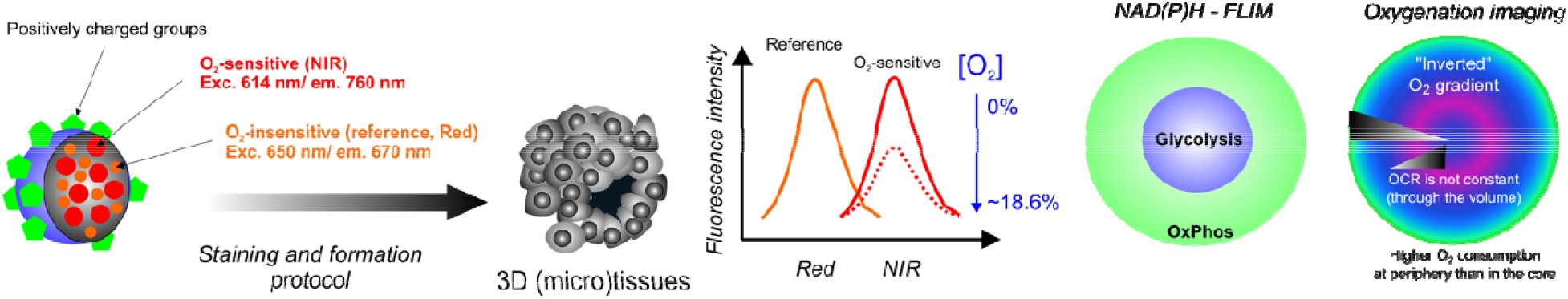

---

Oxygen is a key metabolite for cell metabolism and energy production in the form of adenosine triphosphate (ATP), *via* oxidative phosphorylation. Under physiological *normoxia*, O_2_ supply and consumption are balanced, while in *hypoxia* (or oxygen-deprived conditions, compared to physiological tissue level) O_2_ concentration is below the norm. Different tissues often show different oxygen requirements depending on their function, metabolite consumption and vascular oxygen supply, often ranging from 2 to 9% O_2_ (14-65 mmHg).^1^ Hypoxia results in diminished mitochondrial activity, overproduction of reactive oxygen (ROS) and nitrogen (RNS)^2^ species, activation of hypoxia-inducible factor (HIF)-dependent pathways, and ultimately cell death.^3,4^ Therefore, heterogeneity of cell and tissue oxygenation plays an important role in tissue development and homeostasis. Hypoxia is associated with several diseases such as ischemia,^5,6^ cancer, ^7^ cardiovascular disorders,^8^ inflammatory,^9,10^ and infectious diseases.^11–13^ In solid tumours, uncontrolled cell proliferation, and abnormal vascularisation lead to hypoxia,^14^ resulting in poor prognosis, increased cancer cell survival, metabolic switch from oxidative phosphorylation towards aerobic glycolysis,^15^ increased cell migration,^16^ and resistance to therapy.^17^ 3D tumour spheroids have been widely used to study oxygenation gradients and hypoxia *in vitro,* as promising models to bridge the gap between monolayer cultures and intravital experiments, mimicking organ-specific tissue architecture and microenvironment.^18–21^ Typically, tumour spheroids larger than 300-500 µm in diameter are expected to consist of three concentric layers:^22–24^ the proliferating, quiescent, and necrotic cores, due to nutrients and oxygen diffusion limits concomitant with accumulation of waste products, lactate, and decreasing pH.^25,26^

Traditional methods for (intra)cellular oxygen measurements and oxygen consumption rates (OCR) in such 3D models as spheroids, neurospheres, organoids, and (micro)scaffold-grown structures include microelectrodes,^27^ redox-sensitive nitroimidazole derivatives,^28^ indirect staining with antibodies and hypoxia markers (such as HIF-1α),^29^ genetically encoded fluorescent reporters,^30,31^ organ-on-a-chip devices coupled with solid-state sensors, ^32,33^ optical-based multi-well plate systems, ^34,35^ and optical methods using fluorescent^36–38^ and phosphorescent^39–44^ probes. Optical sensing of molecular oxygen (O_2_) has gained significant interest as it allows for live monitoring of cell metabolism, OCR, and oxygen gradient in a direct, non-invasive, non-chemical, and highly sensitive manner with broad possibilities for multiplexing.^45^ O_2_ indicators are designed based on the phosphorescence quenching phenomenon of the macrocyclic metal complexes, such as Pt(II),^46,47^ Pd(II)^48^ metalloporphyrins and Ru(II) polypyridyl complexes,^49^ where O_2_ interacts with molecules in the triplet-excited states causing a non-emissive deactivation of the phosphor, resulting in a reduction in luminescence.^50^ Thus, higher O_2_ levels lead to lower phosphorescence intensity and shorter decay times (lifetimes), while lower O_2_ levels lead to higher phosphorescence intensities and longer lifetimes.^51^

Phosphorescent probes are commonly designed to be shielded from undesirable microenvironmental effects by either adding a physical barrier, *via* chemical modification with protective chemical groups,^52–54^ or encapsulated into biocompatible and oxygen-permeable nanoparticles (NP).^45^ Chemical modification can help improve cell penetration and biocompatibility, while polymer encapsulation finetunes the quenching sensitivity and protects indicators from local environmental influences (*e.g.* low pH) and interactions with biological components (*e.g.* albumin).^55^ MitoImage^TM^-NanO2 was one of the first-generation NP nanosensors, containing the hydrophobic phosphorescent Pt(II)-tetrakis(pentafluorophenyl)porphyrin dye (PtPFPP) in the cationic RL-100 polymer, allowing intracellular O_2_ measurement in phosphorescence lifetime imaging microscopy (PLIM) and time-resolved fluorescence plate reader of 2D and 3D cell cultures.^56^ However, the microscopy set-up, the intensity of the excitation light, excitation time, scattering, and sensitivity of the photodetector, can influence the phosphorescence intensity of the O_2_ indicators. Ratiometric analysis with the help of O_2_-sensitive and added insensitive indicators can reduce these effects through the use of internal O_2_ calibration,^46^ such as with MitoImage^TM^ MM2 probe,^47^ conjugated polymer NPs,^43,57^ ratiometric FRET NPs,^58,59^ polymer dots,^60^ and negatively charged PMMA-MA NPs.^61^

Surprisingly, there has been little attention to the use of far-red and near-infrared (NIR) dyes in O_2_ imaging so far.^62–65^ Using such structures with long-wave excitation and emission would provide decreased phototoxicity, ensure better filtering of the autofluorescence^66^ and provide deeper light penetration across the volume of multicellular spheroids. While some progress has been achieved with the dye- and fluorescent protein-based structures, their use in ratiometric measurements in imaging 3D microtissues is still rare.^41,67–70^ Here, we demonstrate the red/near infrared-emitting nanoparticle probes, which provide non-toxic, stable, and cell line-dependent staining, and can be used for long-term monitoring of rapid changes in oxygenation in multicellular spheroids. We also validate the detection of observed gradients with the label-free two-photon FLIM microscopy of NAD(P)H. Collectively, the presented approach should help to standardise studies probing hypoxia in 3D tissue models.

## Results and discussion

### Design and spectral characterization of the MMIR probes

To ensure more efficient O_2_-sensing with the help of far-red and NIR dyes, we designed MMIR sensor probes by nanoprecipitation technique^71^ in which the O_2_-sensitive phosphorescent reporter dye, PtTPTBPF^72,73^ (exc. 620 nm, em. 760 nm) and the reference O_2_-insensitive fluorophore aza-BODIPY^74^ (exc. 650 nm, em. 675 nm) are impregnated in the nanoparticle-forming cationic polymer RL-100 (Eudragit) (Fig. 1A, B). The dyes were selected according to the following criteria: (i) hydrophobicity to ensure no leaching and stability in the nanoparticles; (ii) efficient excitation in the red part of the spectrum for better compatibility with biological probes; (iii) significantly different position of the emission maxima for ratiometric read-out; (iv) absence of spectral overlap between the emission of the oxygen indicator and the absorption of the reference dye and between the emission of the reference dye and the absorption of the oxygen indicator in order to avoid Förster resonance energy transfer (FRET). Fig. 1B-D shows that these conditions are fulfilled for the selected pair of dyes. It should be noted that PtTPTBPF is also commercially available, and the aza-BODIPY dye can be prepared with moderate synthetic effort.^74^ As can be seen from Fig. S3 and S4, the particles can be prepared in a reproducible manner. Not surprisingly, the emission ratio is significantly influenced by the ratio of both emitters so that resulting spectral properties can be affected by possible errors (during dye weighing and pipetting of stock solutions), as was observed for batch 1 of MMIR beads. To minimize such fluctuations, it is therefore advisable to produce larger batches of the nanoparticles (i.e. >> 50 mg, the amount used in this work). UV-VIS spectroscopy appears to give a simple way to screen for potential flaws in particle preparation (Fig. S3). Initial screening was performed with different dye ratios (1:1, 1: 0.5, and 0.5:1) to select the most optimal material with respect to the available microscope detector. Although all further experiments were performed with 1:1 ratio of the dyes (both 1% wt with respect to the polymer) it is easily possible to adjust the ratio for other equipment if necessary. The particles were found to be excellently suitable for storage (4 °C) over a prolonged period of time (months and even years) without noticeable aggregation or increase in turbidity (Fig. S5).

**Figure 1:**
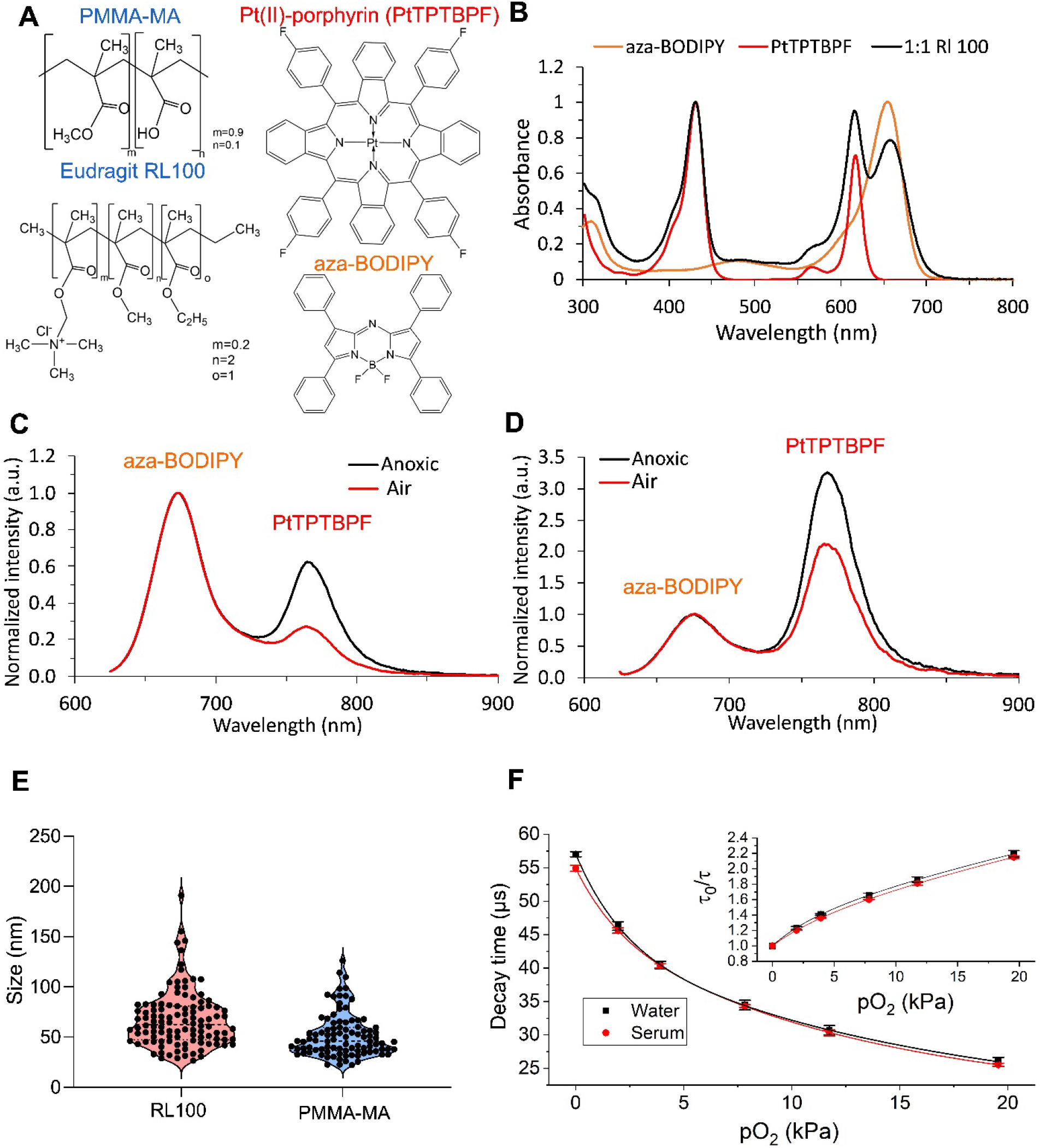
Chemical and spectral characterisation of MMIR O_2_ probes. A: Chemical structures of the reference dye (aza-BODIPY) and O_2_-sensitive (PtTPTBPF) co-precipitated in either PMMA-MA or Eudragit RL100 nanoparticles. B: Absorption spectra of the resulting MMIR probes (1:1 dye ratio) as well as the absorption spectra of both dyes in toluene solution. C: Emission spectra of MMIR probe (1:1 ratio of PtTPTBPF to aza-BODIPY) in anoxic and air-saturated water (λ_exc_ 615 nm). D: Emission spectra of MMIR-probe (1:1 ratio of PtTPTBPF to aza-BODIPY) in anoxic and air-saturated water (λ_exc_ 615 nm). E: Size distribution of the RL100 (N= 230) and PMMA-MA (N=190) nanoparticles measured using a transmission electron microscope. F: O_2_ response and Stern-Volmer relationship (inset) for the phosphorescence lifetime of MMIR1, (23 °C) in water (black) and 10% serum (red). The data represent an average for three different batches of the material.

Previous work ^61^ showed the anionic poly(methyl methacrylate-*co*-methacrylic acid) (PMMA-MA) nanoparticle PA2 showed better staining of multiple neural cell lines and 3D tissue models and had fewer precipitation issues compared to the cationic nanoparticles.^40^ Therefore, we also produced such probe termed “MMIR-“, containing 1 wt% aza-BODIPY and 1 wt% PtTPTBPF (Fig. 1A). This material also showed ratiometric oxygen sensing capability (Fig. 1D) but the sensitivity to oxygen was found to be much lower than in case of MMIR1 probe which is explained by higher permeability of the polymer matrix for the latter. Despite the identical ratio of both dyes in MMIR1 and MMIR-materials (Fig. S3) the emission spectra of the latter indicate a significantly stronger contribution from the oxygen indicator compared to the reference dye (Fig. S4). Excitation of both dispersions used in the same concentration and showing identical absorption at the excitation wavelength revealed comparable emission intensities for PtTPTBPF in MMIR1 and MMIR-(Fig. S6). However, the fluorescence of aza-BODIPY appeared to be significantly quenched (∼ 10 fold) in MMIR-compared to MMIR1. Measurement of fluorescence decay times revealed a similar degree of quenching (Fig. S7) with decay times of 4.3 and ∼0.5 ns in MMIR1 and MMIR-, respectively. This may be due to poor compatibility of the reference dye with PMMA-MA matrix resulting in aggregation of the fluorophore since the decay time for the nanoparticles that contained only the aza-BODIPY dye was also found to be short (∼1.6 ns). In contrast to the reference dye, PtTPTBPF was found to be much better compatible with the PMMA-MA matrix. In fact, the measurement of phosphorescence decay times of PtTPTBPF in MMIR1 and MMIR-materials (Fig. S8 and S9) did not show such a drastic quenching (decay times in deoxygenated conditions of 57 and 39 µs, respectively).

The morphology and size of both MMIR nanoparticle types were analysed by transmission electron microscope (TEM) and dynamic light scattering (nanoparticle tracking analysis, NTA). Interestingly, NTA revealed (Fig. S10A) that PMMA-MA displayed high reproducibility of production and size distribution of ∼64 nm, while RL100-based nanoparticles did not efficiently scatter the light and their size was seemingly in the range of <60 nm, comparable with previously reported size measurements for MM2 (70 nm) and PA2 (95 nm) probes^61,74^. TEM showed similar results (∼68 nm for RL-100 and ∼52 nm for PMMA-MA) but with less regular shapes, which can be explained by the aggregation of these polymers when present in a dried form (Fig. 1E, S10B). Thus, PMMA-MA in dried form (TEM) displayed smaller size, which could be explained by the differences in the size of the hydration shell and scattering efficiency. Next to the possibility of fluorescence intensity-based ratiometric analysis using common widefield and confocal fluorescence microscopy, the nanoparticles should also display O_2_-sensitive phosphorescence lifetime changes measurable by phosphorescence lifetime imaging microscopy (PLIM). We performed calibration of MMIR1 in water and 10% serum (mimicking cell environment) and found a minor effect of serum, slightly reducing of the luminescence decay times. This can be explained by the fact that some population of the dye is still exposed to the ’’external’’ microenvironment and is not fully protected, due to the nature of the nanoprecipitation technique. Nevertheless, the effect was minor (an average of 4%) and we concluded that nanoparticles provide good protection against interferences from serum and other biological components (Fig. 1F). These measurements also indicated high reproducibility of the oxygen sensing properties for different batches of the material even if the ratio of both emitters is slightly varied. For the subsequent experiments, we decided to focus on ratiometric semi-quantitative detection, as the most widely available and affordable microscopy readout.

### Cell staining, localization, and effects on cell viability of MMIR probes

First, a selection of the produced nanoparticles was made based on their staining efficiency after overnight staining (17 h) with a ‘standard’ concentration for nanoparticles (5 µg/ml) on adherent cell monolayer cultures, including human colon cancer cells (HCT116), human Dental Pulp Stem Cells (hDSPC), and Human Umbilical Vein Endothelial Cells (HUVEC) (Fig. S11).

Positively charged particles displayed an overall higher fluorescence intensity compared to the negatively charged ones. As expected, MMIR-showed fewer probe aggregation when in contact with cells, compared to the cationic MMIR1 (0.5:1 and 1:0.5 ratios) (Fig. S11, A-B).

Subsequently, we selected MMIR- and MMIR1 for the following tests on effects on cell staining, localization, cell viability, and experiments with 3D cultures.

We studied nanoparticle concentration-dependent cell staining using overnight incubation with a range of concentrations (0 - 20 µg/ml) (Fig. 2A). This helped us to find 5 µg/ml as sufficient staining concentration for MMIR1 and 20 µg/ml for MMIR-. Next, we looked at the kinetics of cell staining and found that NPs showed intracellular uptake after 4 hours of incubation, reaching maximal signals after 17 h for MMIR1 (Fig. 2B). Longer incubation led to noticeable precipitation outside the cells. Interestingly, MMIR-showed a steady increase in signal even after 24 hours.

**Figure 2:**
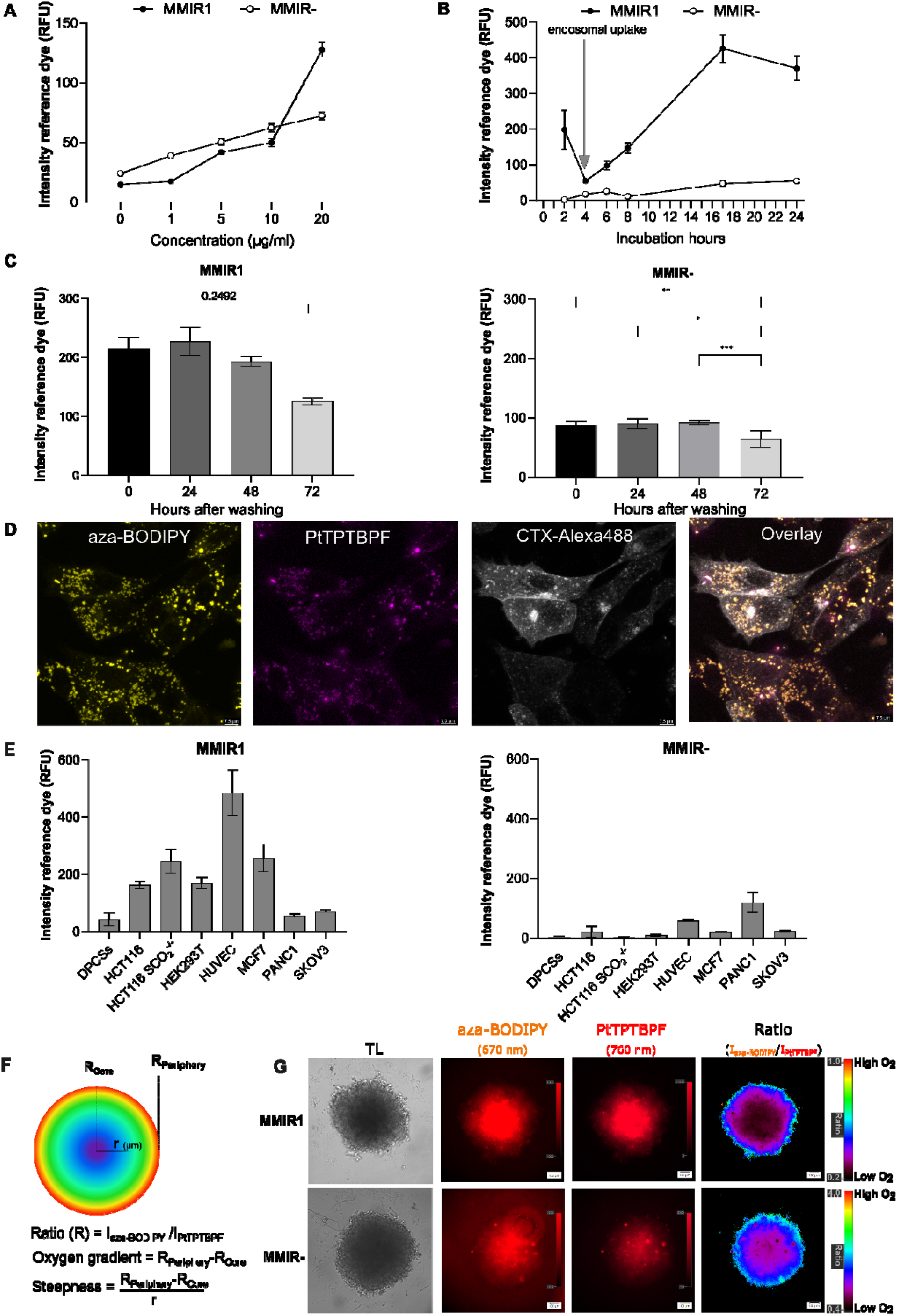
Cell staining and localisation of MMIRs in live cells. A: Concentration dependence of intracellular uptake of MMIR probes. HCT116 cells were incubated with nanoparticles (0-20 µg/ml, 18 h), washed, and quantified on a fluorescence microscope. Data shows the average of 3 repeats (no background subtraction) ± SEM. B: Time-dependent staining of HCT116 cells with MMIR nanoparticles. Cells were incubated MMIR (0-24 h, 5 µg/ml for MMIR1 or 20 µg/ml for MMIR-). Data shows the average of 5 repeats (with background subtraction) ± SEM. C: Retention of fluorescent signals (reference dye) of MMIR1 (5 µg/ml, 17 h) and MMIR-(20 µg/ml, 17 h) in HCT116 cells after staining. The data shown is an average of 4 repeats (with background subtraction) ± SEM. *: P= 0.0158, **: P= 0.0091, ***: P= 0.0002. D: Intracellular localization of MMIR1 after overnight incubation with HCT116 cells (5 µg/ml, 17 h) shows endo-and lysosomal localization. Cells were co-stained with Cholera Toxin-Alexa Fluor 488 (2 µM, 1 h) and imaged using confocal microscopy. Scale bar is 7.5 µm. E: Cell line-dependent uptake of the MMIR1 and MMIR-probes ( 5 µg/ml, 17 h), based on fluorescence microscopy experiments. Data shows the average of 3 replicates (with background subtraction) ± SEM. F: Quantitative formulas for statistical analysis. R= Ratio, I= intensity, r= radius. G: Ratiometric imaging of the oxygen gradient using MMIR1 and MMIR-probes in live SKOV3 cell spheroids with the shown transmission light (TL), sensitive, and reference dye intensity images. Scale bar is 100 μm.

To investigate how long nanoparticles could remain inside the cells, we performed a ’leakage’ experiment over 72 hours (Fig. 2C). The MMIR1 remained in the cells with a non-significant reduction of reference dye intensity for up to 72 h (P=0.2492), while the MMIR-probe had a significant reduction of signals after 72 h (P= 0.0091). These differences can be explained by probe dilution upon cell division and different cell entry mechanisms, dependent on the type of nanoparticles. With spheroid cultures, we found that keeping MMIR1 in the growth media resulted in long-term retention for up to 26 days (Fig. S12).

Cells take up most commonly available O_2_ sensing nanoparticles through the mixed endocytosis mechanisms ^56,75^ leading to predominantly endo- and lysosomal localisation. This was confirmed with our MMIR probes by incubating live HCT116 cells overnight with MMIR1 (5 µg/ml) and co-stained with green organelle-specific marker dyes (Fig. 2D, Fig. S13). The mixed endocytosis mechanism was further confirmed by the cell-line dependent staining efficiency for both nanoparticle types (Fig. 2E). We found that MMIR1 efficiently stained most cancer and non-cancer cell lines, while for MMIR-more cell-specific behaviour was noted (Fig. S14).

Considering the rate and mechanism of cell entry for MMIR probes, we could stain 3D multicellular spheroids models by adding nanoparticles during spheroid formation or forming spheroids from already pre-stained cells (Fig. S15). This method ensures improved distribution of nanoparticles through the volume of spheroids of different sizes and different cell types, including HCT116, DPSC, MDA-MB-231, and SKOV3. Ratiometric analysis ensures the visualisation of the oxygen gradients in the spheroids, through the normalisation of the fluorescence intensity of the reference by the sensitive dye signal. Previously,^63^ we introduced the parameters “oxygen gradient” (R_periphery_ - R_core_) and “steepness” (ΔRatio/spheroid radius), hereby making quantitative measurements possible (Fig. 2F). We found that ovarian adenocarcinoma SKOV3 cells internalized both MMIR nanoparticle types with similar efficiency (Fig. 2E), and SKOV3 spheroids displayed comparable oxygenation gradients (Fig. 2G).

Potential cytotoxic effects of MMIR nanoparticles were investigated with both 2D monolayer and 3D spheroid cultures. We looked first at the membrane integrity assay with Sytox Green dye and saw no statistically significant effects on cell viability (Fig. S16, A). Furthermore, we measured total cell ATP levels and saw no effect for the concentration range of MMIR1 up to 50 µg/ml for HCT116 cells (Fig. S16, B). This was also confirmed by the less direct MTS viability assay method (Fig. S17). MMIR1 also did not influence cellular ATP levels upon activation of mitochondria with uncoupler FCCP and inhibitor Oligomycin, confirming no effect on oxidative phosphorylation (Fig. S16C). To confirm the low potential cytotoxicity of the MMIR1 to the cells, we tested its effect at the highest staining concentration (50 µg/ml) on the broader panel of cell lines, which we used for cell staining experiments (Fig. S16D): no statistically significant effect on the cell function was observed. For HCT116 cells, we extended our tests to 3D cell culture and looked at the effects of staining with MMIR1 on resting ATP in spheroids using the CellTiter-Glo 3D viability Assay (Fig. S16E). Finally, we also looked at the effects on the distribution of fluorescence lifetimes of endogenous NAD(P)H in MMIR1-stained spheroids *via* two-photon FLIM: no effects on cellular redox were observed (Fig. S18).

### Evaluation of MMIR1 nanoparticles in monitoring oxygenation in multicellular spheroids

To confirm the photostability of MMIR1 fluorescent signals in stained spheroids, we performed repeated illumination (10 cycles) on a conventional LED-based fluorescence microscope: we observed less than a 5% drop in initial intensity ratio, in agreement with the high photostability reported previously for other nanosensors,^56,75^ compared to reference tetramethylrhodamine methyl ester (TMRM, P<0.001) (Fig. S19A). Thus, both MMIR probes allow for dynamic real-time study of rapid respiratory responses to mitochondrial uncouplers, activators, and inhibitors of the electron-transport chain. We found that in HCT116 spheroids, FCCP only displayed a mild uncoupling effect at the spheroid core within 2 minutes of the drug addition. However, a slight increase in the ratio was observed at the spheroid periphery. This is in agreement with previously reported metabolic features of spheroids, which require at least 10 minutes to display a clear uncoupling effect.^61–63^ Rotenone strongly inhibited mitochondrial respiration causing spheroids reoxygenation, leading to an increased ratio % at both spheroid’s periphery and core within ∼30 seconds after stimulation (Fig. S19B).

Additionally, responses to O_2_ changes were demonstrated in oxygenated (respiratory inhibition by antimycin A and rotenone) and deoxygenated HCT116 spheroids. The ratiometric intensity changes at the periphery were significant (P<0.0001) (Fig. S19C). This illustrates the applicability of the ratiometric readout with MMIR probes for quantitative monitoring of rapid changes in oxygenation in spheroids.

### Factors affecting the shape of O_2_ gradients in spheroids

The common mechanism of spheroid formation is based on self-assembly in a non-adherent environment, leading to compactization by E-cadherin.^76^ Multiple techniques such as micro-patterned surfaces, hanging drop, spinner flasks, use of non-adhesive surfaces, or laser-based 3D bioprinting can produce spheroids of different sizes.^77,78^ Even though 3D models gain significant interest compared to 2D cultures, their reproducibility and high variability remain a problem.^40,44,79^ Such factors as the nutrient composition of culture media,^79–81^ spheroid formation method,^82^ and the viscosity of the extracellular fluid^83,84^ can contribute to changes in cell viability, differentiation capacity, and response to (bio)chemical signals and thus result in heterogeneity of the grown spheroids and organoids.^85–87^

Multicellular tumour spheroids are considered to be a “simple” model, developing ‘classical’ size-dependent diffusion gradients of nutrients, ATP, waste, and molecular oxygen (O_2_), with three concentric structures representing proliferating, quiescent, and necrotic zones.^22–24^ In the necrotic core, limited O_2_ diffusion creates a hypoxic or anoxic region, which can be visualized using our O_2_ probe. Thus, live HCT116 spheroids showed size-dependent changes in their oxygenation (Fig. 3A, Fig. S20A) at the core and periphery (Fig. S20B-C). However, the oxygenation steepness, or changes in oxygen gradient per µm, were not significantly size-dependent (Fig. S20D).

**Figure 3:**
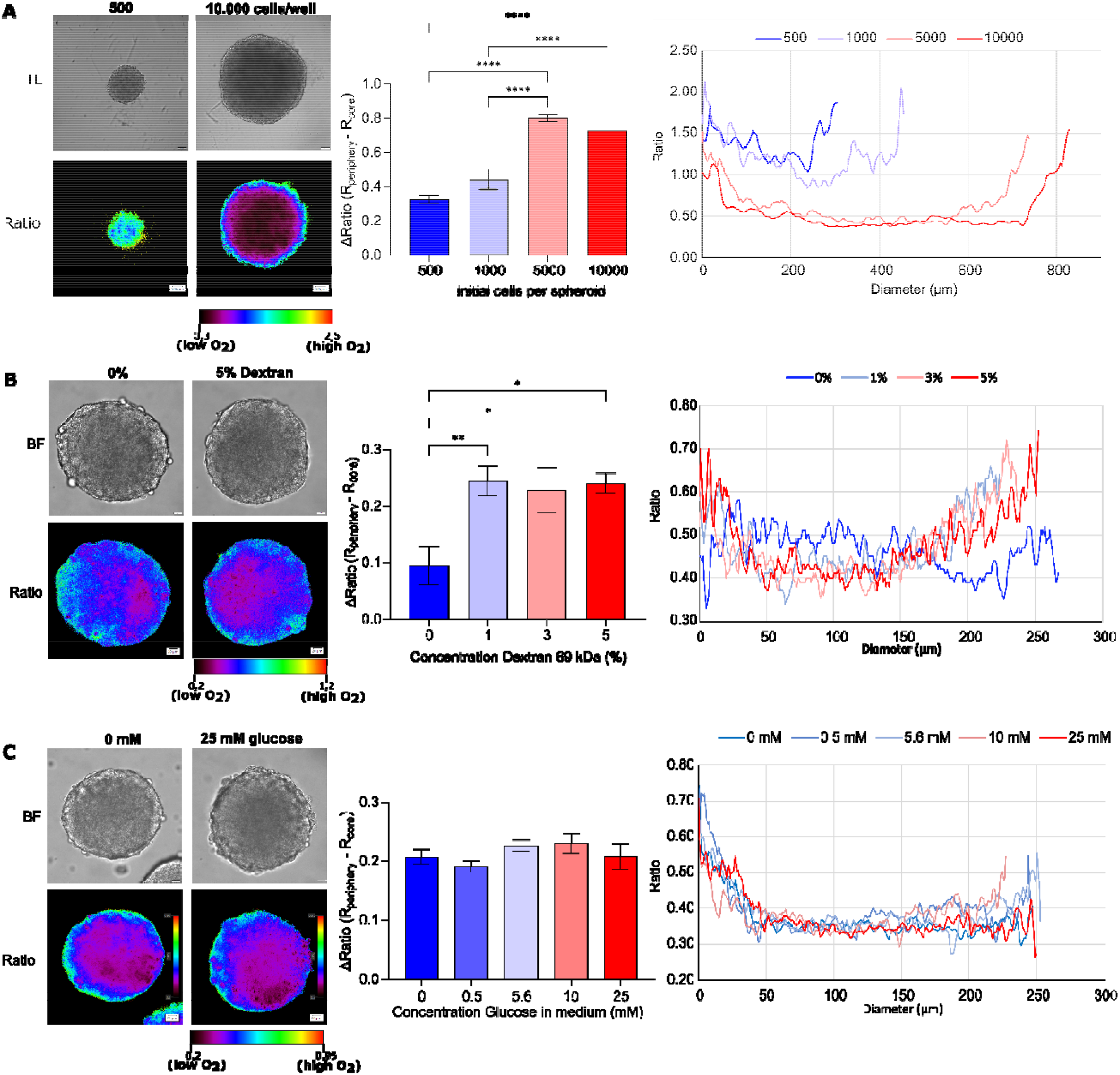
Oxygenation gradients are influenced by size, extracellular viscosity, and glucose concentration. A: Size-dependent oxygenation of live HCT116 spheroids. HCT116 spheroids formed on a Lipidure^®^-coated plate from 500, 1,000, 5,000, and 10,000 cells per well (5 days). Results show the average ± standard error of 6 spheroids. Scale bar is 100 μm. B: Increased cell media viscosity results in the formation of a more hypoxic core. HCT116 spheroids formed using the agarose micromold method were adapted to imaging media containing 0-5 w/w% dextran (0.77-2.25 cP) for 4 hours before imaging. Scale bar is 20 µm. Results show the average ± standard error of 9 spheroids. C: Effect of glucose concentration on oxygenation of live HCT116 spheroids. Spheroids were formed using the agarose micromold method and adapted to imaging media containing 0-25 mM glucose. Scale bar is 20 µm. Results show the average ± standard error of 9-14 spheroids.

Extracellular fluid viscosity has been recently shown to increase the motility of various 2D breast cancer cell lines (*e.g.* MDA-MB-231) and cell dissemination in 3D tumour spheroids.^83,84^ However, increased viscosity should also lower the oxygen delivery and result in a more hypoxic environment, which was not studied by Bera *et al*.^84^ To test this, we exposed live HCT116 spheroids to the growth media of higher viscosity, by supplementing with 0-5 w/w% dextran (0.77 cP to 2.5 cP, 4 h) (Fig. S21). We found that spheroids incubated under higher viscosity compared to growth media (0.77 cP), showed a significant increase in oxygenation gradient (Fig. 3B), produced a more hypoxic core (Fig. S21C), and significantly increased steepness of oxygenation (core to periphery) (Fig. S21D). Thus, an increased extracellular viscosity can also contribute to the “hypoxic cell priming” and affect the metastatic cell dissemination.^88^

We also reasoned that the glucose concentration in the medium could influence oxygenation in spheroids, similar to our previous findings with small intestinal organoids^89^ and indirect evidence from the literature.^79^ Thus, we looked at the effect of acute change of the medium glucose content (0-25 mM, 4 h) on the oxygenation of the HCT116 spheroids (Fig. S22). Interestingly, while some changes in oxygenation and steepness (Fig. S22D) were seen, they were not statistically significant (Fig. 3C). This means that short-term changes (2-4 h) in the growth medium may not significantly affect spheroid oxygenation (and potentially cell metabolism and viability), in contrast to longer-term exposure times (5-7 days) affecting cell death as seen by Baye and coworkers.^90^

The spheroid formation method can greatly influence spheroids variability, size, and shape, resulting in cell density, and drug sensitivity.^62,79,82,91^ Therefore, we investigated how the different high throughput (SphericalPlate 5D^®^, self-made micromolds, and Microtissue^®^ molds) and the ‘medium throughput’ low attachment (Biofloat^TM^ and Lipidure^®^-coated 96-well plates) methods affect the oxygenation, morphology, and viability of HCT116 spheroids (Fig. 4).

**Figure 4:**
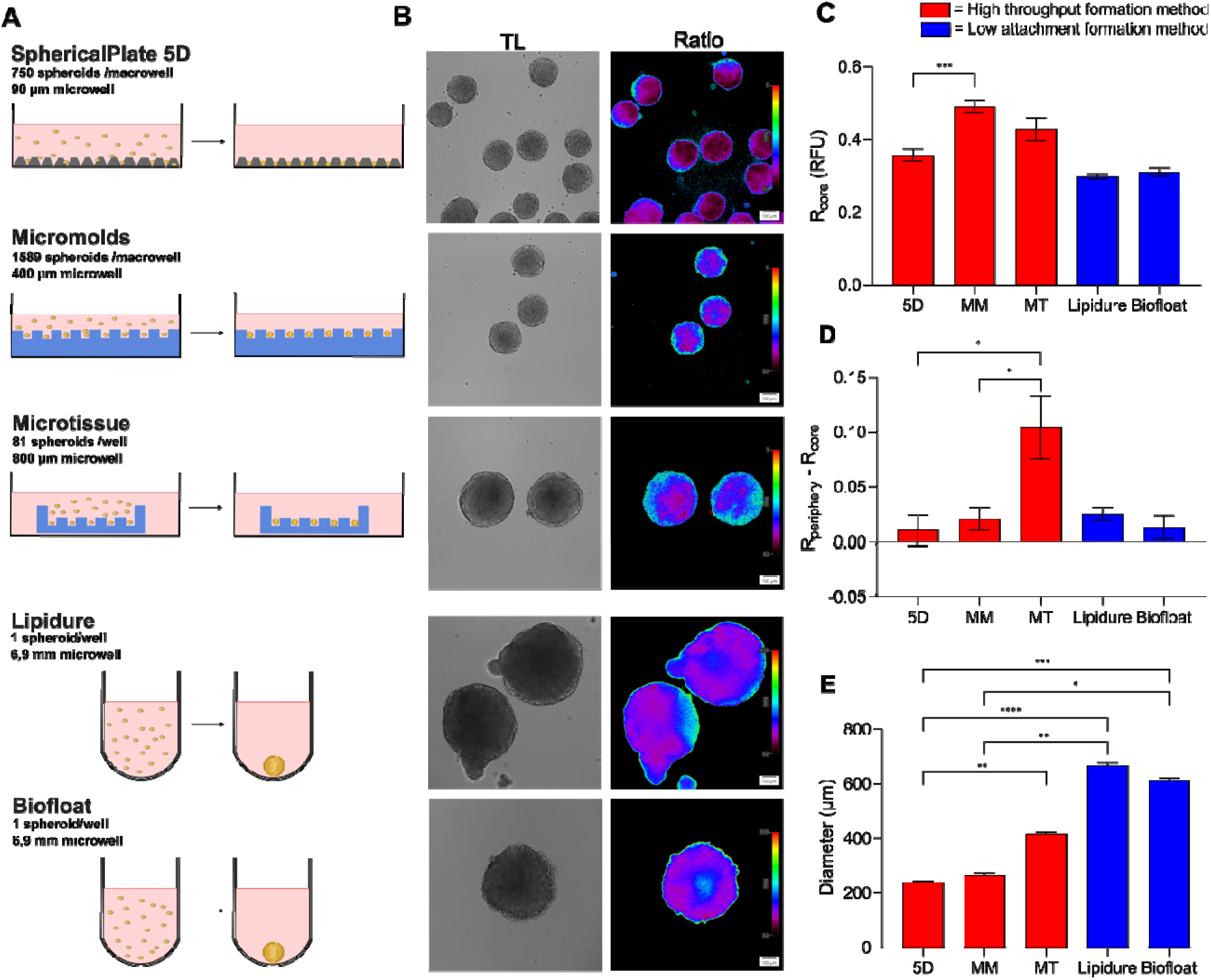
Spheroid formation methods affect morphology, size, viability, and oxygenation. A: High-throughput methods such as the self-produced micromolds and the Microtissue^®^ molds use stamps to make multiple microwells in agarose (blue), while the SphericalPlate 5D^®^ has integrated patented microwell in the plate. Low-attachment plates such as Lipidure^®^ (Amsbio) and Biofloat^TM^ (Sarstedt) use a non-adherent coating inhibiting cell-surface adhesion and promoting cell self-aggregation. B: Oxygenation in HCT116 spheroids (initial seeding amount of 500 cells, 20 µg/ml, MMIR1, 6 days) produced by different methods showed more oxygenated spheroids with the MicroTissue and self-made Micromolds than the 5D SphericalPlate^®^. C: MMIR1 ratio measurements at the core. D: Oxygenation gradients. E: Spheroids size. Results show the average ± standard error of 6-16 spheroids. 5D= SphericalPlate 5D^®^, MM= Micromold method, MT= MicroTissue^®^ method.

Chosen high throughput methods use either integrated grids or microwells in agarose to produce multiple spheroids, while a non-adhesive coating is used in the low-attachment plates hereby stimulating self-assembly (Fig. 4A). Surprisingly, spheroids formed on low attachment plates were significantly larger, with 667.5 ± 22.65 µm (Lipidure^®^) and 612 ± 25.33 µm (Biofloat^TM^), compared to 5D plates (239.6 ± 9.06 µm), Micromolds (265.3 ± 37.45 µm) and Microtissue^®^ (417.8 ± 15.62 µm) (Fig. 4E). Low attachment methods resulted in clear development of necrotic core seen with propidium iodide staining (Fig. S23A). This difference in cell viability is expected to attenuate oxygen diffusion through the media, which in turn can be affected by O_2_ partial pressure, O_2_ consumption rates, height of the culture to cell distance, temperature, and O_2_ solubility.^92,93^ Furthermore, nutrients get depleted and waste products accumulate faster with a larger number of spheroids within the same volume and therefore require more frequent media exchange.

The SphericalPlate 5D^®^ spheroids showed lower oxygenation at the periphery (Fig. S23B) and core (Fig. 4C) compared to the Microtissue^®^ (respectively P= 0.013 and P>0.9999) and the self-made micromolds (respectively P= 0.0007 and P= 0.0010). This can be explained by the greater distance from the spheroid to the surface, leading to slower O_2_ delivery in the static culture. This hypothesis is valid as oxygenation in Biofloat^TM^ and Lipidure^®^ low attachment plates are statistically similar. As we have seen previously, size differences between the high throughput methods cause significant differences in overall spheroids oxygenation (Fig. 4D) but not in their steepness (Fig. S23C). Most surprisingly, the low-attachment spheroids displayed increased oxygenation in their core, depending on their size and time after formation (Fig. S24).

Thus, the MMIR1 probe demonstrated utility in monitoring cell oxygenation, in dependence on spheroid size, extracellular medium viscosity, and composition. In addition, we observed that the different spheroid formation methods can lead to drastic differences in spheroid composition and viability, agreeing with earlier studies and meta-analysis.^79^

### Inverted O_2_ gradient in spheroids

With some spheroid preparations, produced with *e.g.* low-attachment methods (Fig. 4B) we encountered an unusual “inverted gradient” distribution of hypoxia, when O_2_ in the core was somewhat higher than at the periphery. Interestingly, this phenomenon was observed with spheroids produced from cell lines of completely different origins, including human dental pulp stem cells (DPSCs, stem cell line) and colon cancer HCT116 cells (Fig. 5 and Fig. S25). Thus, with hDPSCs, small size spheroids showed ‘classical’ O_2_ distribution, while upon growing to larger sizes or when seeded with higher cell numbers (160 μm size, Fig. 5A-C), the shape of the gradient changed. A similar situation was observed with HCT116 cells, although with spheroids of much larger size (Fig. 5D-F).

**Figure 5:**
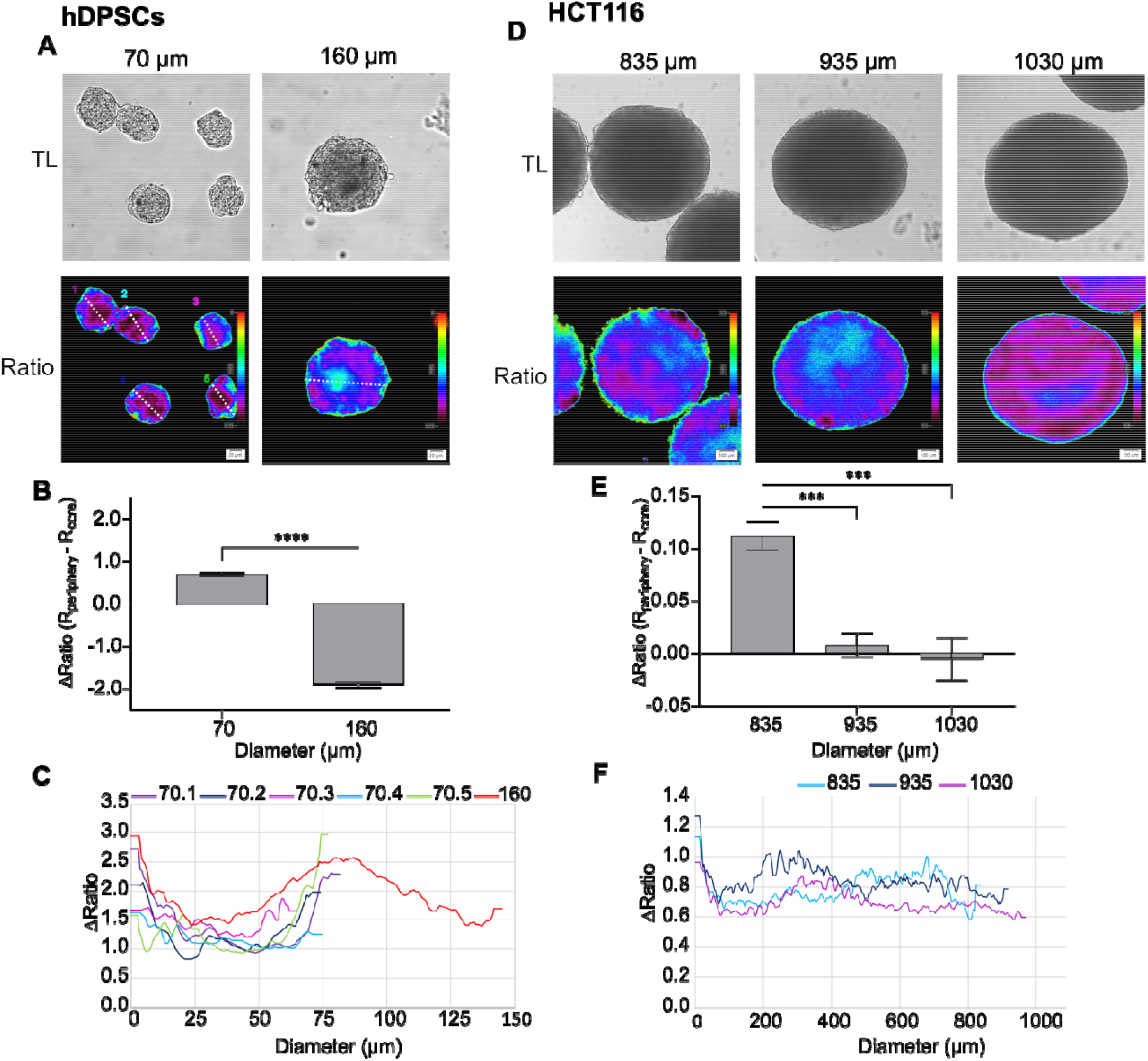
Inverted oxygen gradient in non-tumour and tumour spheroids. A: hDPSCs spheroids formed using agarose micromolds in 2 sizes (70 µm and 160 µm) for 8 days by respectively seeding 3×10^5^ and 1.3×10^6^ cells per macromold. Scale bar is 20 µm. B: Oxygenation gradient is measured by subtraction of the R_core_ from R_Periphery_. Results show the average ± standard error of 30-33 spheroids. ****: p<0.0001. C: Ratiometric line profiles show increased oxygenation at the core in bigger spheroids. D: HCT116 spheroids (P. Hwang, NIH cell line) formed for 7 days on Lipidure^®^-coated plate (Amsbio) by initially seeding 1,000, 10,000, and 20,000 cells per spheroid, resulting in respectively 835, 935, and 1030 µm diameter. Scale bar is 100 µm. E: Oxygenation gradient is measured by subtraction of the R_core_ from R_Periphery_. Results show the average ± standard error of 8-10 spheroids. ***: p<0.005. F: Ratiometric line profiles show an increased oxygenation at spheroids >835 µm.

The observed phenomenon agrees with earlier studies performed by Sutherland and co-workers, that used microelectrode-based O_2_-sensing in mouse epithelial breast cancer EMT6/Ro^94^ and human colon adenocarcinoma Co112 spheroids.^24^ Thus, a reversed correlation between central pO_2_ and glucose supply in large spheroids (>900 µm) was reported. This suggests the potential metabolic adaptation of cells to the different microenvironmental conditions,^94^ occurring in the vicinity of a necrotic core in large-size spheroids (Fig. S26). Other studies using indirect hypoxia labelling with pimonidazole, also reported a circular-shaped hypoxic area adjacent to the necrotic core in T-47D human breast cancer spheroids at day 8.^95^

Furthermore, we investigated the cellular metabolism using the autofluorescence of reduced nicotinamide adenine (phosphate) dinucleotide (NAD(P)H), a coenzyme involved in glycolysis and oxidative phosphorylation.^89,96^ Its fluorescence lifetime is significantly shorter in its free state (∼0.45 ns) compared to the protein-bound and other states.^97^ Using low-attachment and micromold spheroid formation methods and various cell seeding densities, we looked at the development of an inverted oxygen gradient in spheroids from HCT116 cells^98^. First, using oxygenation analysis we confirmed that spheroids from the two different size populations developed two distinct oxygenation profiles: generally hypoxic (’forward gradient‘ spheroids), and with a pronounced ‘inverted‘ gradients (Fig. S27A). Subsequently, we analysed the spheroids with ‘forward’ and ‘inverted’ gradients (identified by oxygen microscopy) *via* two-photon NAD(P)H-FLIM. We observed a heterogeneous population of spheroids with ‘uniform’ NAD(P)H fluorescence lifetime distribution (the most abundant in the group) and with ’structured’ NAD(P)H lifetime distribution (glycolytic core; less abundant) in a group of “small” size spheroids (Figs. 6, 7A, S27 B-C, ST3). The NAD(P)H-distribution structure heterogeneity of the small spheroid population agreed with our oxygenation data, where an appearance of inverted gradients was revealed by O_2_ analysis (Fig. S27A-B).

**Figure 6.**
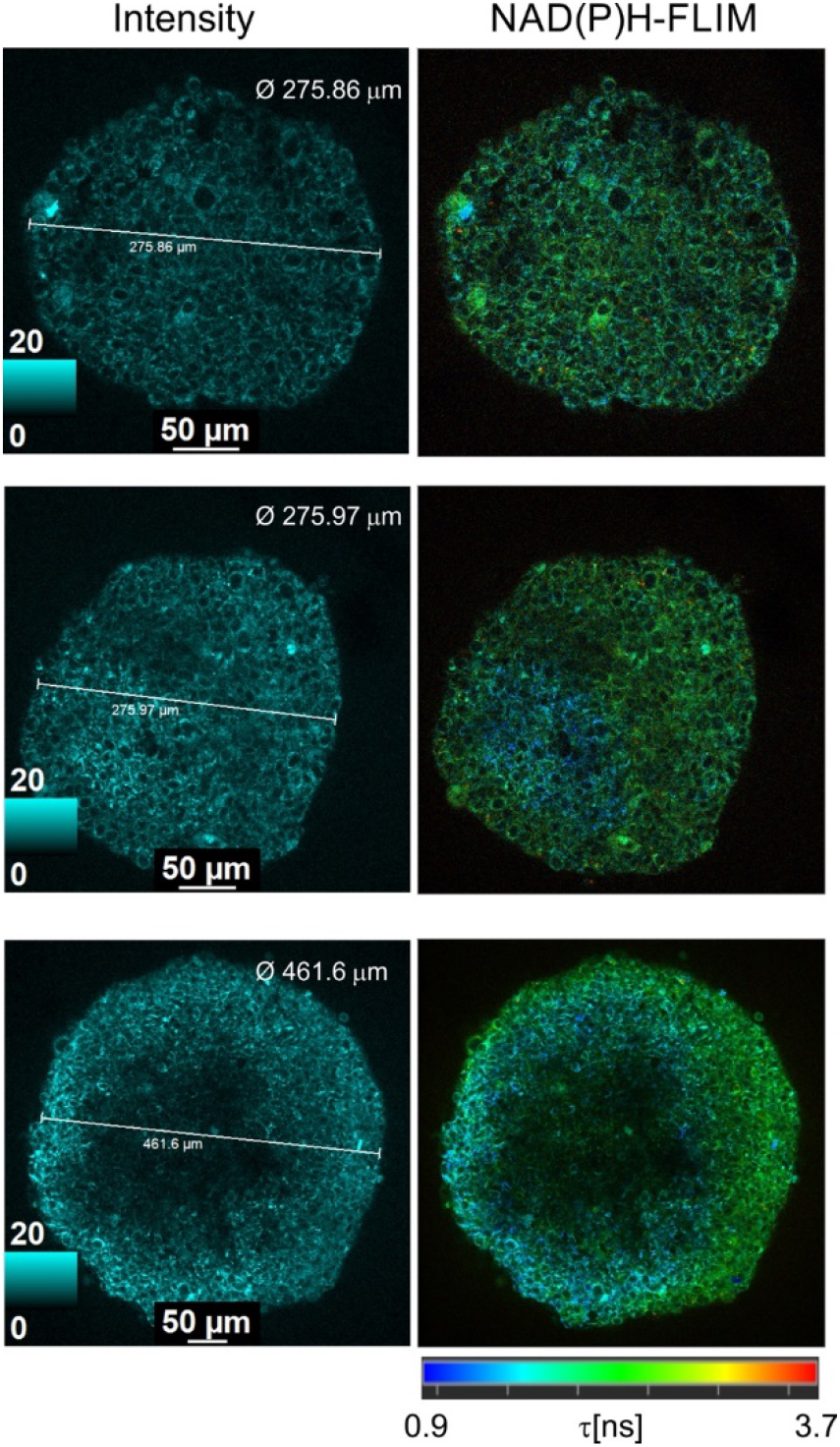
Examples of two-photon NAD(P)H-FLIM microscopy of HCT116 spheroids of different sizes. (increase from top to bottom). Single optical sections (normalised intensity and fast-FLIM images) are shown. Scale bar is 50 μm.

**Figure 7:**
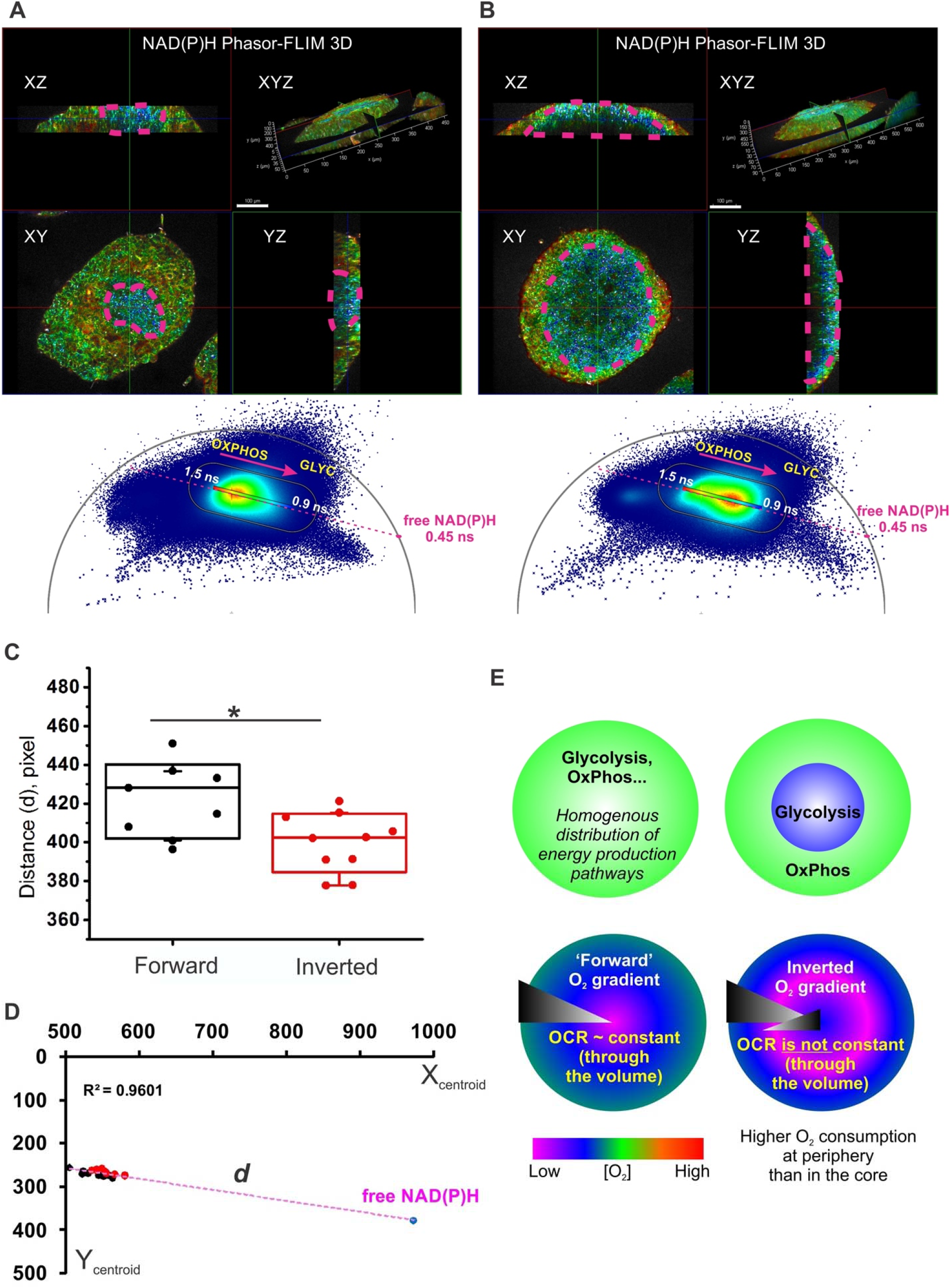
Two-photon FLIM of NAD(P)H reveals a glycolytic switch in HCT116 spheroids with ‘direct’ and ‘inverted’ oxygen gradients. A, B: 3D reconstructions of phasor-FLIM in 3D of early and mature spheroids with ‘inverted’ gradient phenotype. The ‘glycolytic core’ (demarcated by purple dashed line) demonstrated local homogeneity of shorter NAD(P)H lifetime and was surrounded by a zone with longer lifetime distribution. Combined phasor plots of sections from 3D stack of spheroid from the ‘inverted’ gradient group display a characteristic shift towards monoexponential free-NAD(P)H lifetime (0.45 ns) (magenta arrow). Scale bar is 100 µm. C: Comparison of distances (*d*) between the theoretical position of phasor plot of free-NAD(P)H and centroids of free-NAD(P)H phasor plots of spheroids from ‘forward’ and ‘inverted’ gradient groups. Analysed phasor plots corresponded to individual XYZ sections of spheroids imaged within ∼100-200 μm depth from the top of the spheroid. *T*-test (p < 0.05; n ‘forward’ gradient spheroids is 7; n ‘inverted’ gradient spheroids is 9) revealed a significant difference between groups, with the shift of phasor coordinates from ‘inverted’ gradient group spheroids toward the free-NAD(P)H lifetime. Boxes represent standard deviation, whiskers represent 10 and 90 percentiles. D: Linear fitting of phasor plot centroids and free NAD(P)H theoretical coordinates demonstrated accurate linear alignment with R^2^ = 0.96. E: Hypothetical metabolic rearrangements in spheroids during the formation of ‘inverted’ O_2_ gradients.

With the increase in spheroid size, we witnessed the formation of massive glycolytic core regions within spheroids, with clearly shorter (blue) fluorescence lifetimes (Fig. 6 and 7B). Remarkably, some spheroids demonstrated a large glycolytic core, occupying almost half of the spheroid and asymmetrically located close to the spheroid periphery (see example on Fig. 6 bottom panel and Fig. S27C). Similar asymmetry was observed with spheroids having ‘inverted’ oxygenation gradients, allowing to speculate that both methods reveal similar metabolic layers in spheroids and that there is indeed a link between the metabolic stratification and oxygenation profiles within spheroids (Fig. S27A).

We further looked at the spheroid size-dependent distribution of NAD(P)H lifetimes using ‘gold standard’ phasor plot analysis (Fig. S27 and Fig. 7) and demonstrated the appearance of shorter NAD(P)H fluorescence lifetime population in spheroids with ‘inverted’ gradients, reflecting their complex metabolic organisation. The 3D reconstruction of NAD(P)H autofluorescence in spheroids using phasor-FLIM false-colour mask (Fig. 7A-B) illustrates this observation on the example of spheroids with the small (from ‘forward’ gradient spheroid population) and large (from ‘inverted’ gradient spheroid population) glycolytic cores. The spheroid with a large glycolytic core (Fig. 7B) displayed a phasor plot cloud with the appearance of short component lifetimes in comparison to the cloud of spheroid with a small glycolytic core, where the shift towards the long lifetime component was observed. The statistical analysis of phasor plots from the ‘forward’ and ‘inverted’ gradient spheroid groups confirmed the difference between NAD(P)H fluorescence lifetimes, where the position of phasor plot cloud centroids (the linear alignment was detected with R^2^=0.96, Fig.7D) in the ‘inverted’ gradient spheroid group was closer to the theoretical position of monoexponential free-NAD(P)H phasor plot cloud with an average lifetime 0.45 ns (Fig. 7C). This data confirms the dependence of observed fluorescence lifetimes on the metabolic status of cells in HCT116 cells.

Altogether, analysis of forward (normal) O_2_ gradients and NAD(P)H-FLIM revealed the presence of a glycolytic core, which would progressively increase with the size and the spheroid growth time. Interestingly, with increasing spheroid size (> 300-500 μm), this core would also show shorter NAD(P)H intensities, indicating a further decrease in metabolic activity and cell death. Fig. 7E schematically shows a possible explanation of this phenomenon: we hypothesize that spheroids with a ‘homogenous’ distribution of their metabolically active cells (relying either on OxPhos, glycolysis, or a mixture of both) display conventional “direct” O_2_ gradient, determined mainly by diffusion and size. However, spheroids having distinct populations of metabolism will display a more complex oxygenation gradient. This working hypothesis points to the process of spontaneous organised cell metabolic ‘differentiation’, which can occur with some cell lines in the spheroid cultures dynamically. We observed such a phenomenon only with certain cell lines and suspect that the formation of an inverted gradient can be a temporary process, leading to further cell death or further cell differentiation within the ‘mono-culture’.

Rather surprisingly, the observed formation of the glycolytic core in the spheroids and the resulting O_2_ gradient were not reported in the literature previously.^99,100^ This can be explained by the fact that most studies of tumour spheroids focused on these methods separately, often used lower-resolution equipment, and rarely looked at 3D context.^99,101–103^ True, technical limitations of FLIM and PLIM microscopes available on the market^42,104^ rarely allow for such experiments. However, further studies and approaches such as NIR-based nanosensors imaging of oxygenation, are expected to shed more light on this phenomenon.

## Conclusions

In this study, we produced far-red/near-infrared dual emission nanosensor probes MMIR, which show reliable O_2_ quantification in ratio-intensity and decay time readouts. Using the polymeric shell with either cationic or anionic resulting nanoparticles displays cell-specific uptake through mixed-endocytosis with endo- and lysosomal localization, in agreement with our earlier studies with other reference and O_2_-sensing dyes.^47,61^ Within the optimized loading concentrations and incubation times, MMIR probes showed brightness and photostability, ensuring reliable quantification of hypoxia in monolayer and multicellular spheroid cell cultures with no cytotoxic effects (Fig. 1-2). The observed slow cell uptake process (12-16 h) also results in slower release from the cells, which can enable long-term oxygenation monitoring in spheroid cultures for up to 21-26 days.

To illustrate the applicability of the produced nanosensor probe, we focused on ‘semi-quantitative’ ratiometric microscopy, which is widely available, in contrast to the PLIM method.^42,97^ The described approach can be performed on conventional widefield LED-based fluorescence and laser-scanning confocal microscopes and macro-imagers, with a detection sensitivity window of 600-800 nm. While the precise calibration and conversion of fluorescence ratio values into actual O_2_ levels can pose technical challenges for multicellular spheroids and other 3D tissue models^63,70,97,105,106^, it offers real-time and reproducible analysis of ratios, contrasting with the end-point quantification methods^42^.

Collectively, our measurement approach applied to spheroids produced from both stem and cancer cells proved that the oxygenation gradients are not always static. The oxygenation gradients depend upon multiple factors, including spheroid size, extracellular viscosity, and spheroid formation and handling methods. This emphasizes the importance of oxygenation monitoring in studies involving the development and validation of targeted drug/ radio-therapies with cancer cells, tumour organoids, and ‘tumour avatars’.^107,108^ On the other hand, spheroids used as tissue building blocks in 3D printing, bioassembly, and biofabrication, also require monitoring of their viability, cell death, and oxygenation^109,110^. Our ‘discovery’ of the inverted oxygenation gradients with the presented approach and its correlation with autofluorescence label-free two-photon FLIM, highlights the utility of the method, as more affordable, direct, and highly compatible with other multi-parameter micro- and mesoscopy modalities of 3D tissue models.

## Methods

### Materials

Poly(methyl methacrylate-co-methacrylic acid) (PMMA-MA; 10% methacrylic acid, MW ∼100,000 Da, 17913-500) was from Polysciences (USA), organic solvents were obtained from Carl Roth (Austria), Eudragit RL-100 polymer (∼10% of quaternary ammonium groups, MW ∼150,000 Da) was from Degussa (33434-24-2, Degussa, Germany).

Standard cell culture plasticware was from VWR (Belgium). For microscopy, cells were grown or were transferred (as spheroids) onto microscopy dishes (coverglass with No. 1.5 thickness, *e.g.* µ-slide 12-well, Ibidi GmbH, Germany, or equivalent) pre-coated with of 0.07 mg/ml collagen IV/0.03 mg/ml poly-D-Lysine. The following dyes were used: Calcein Green AM (AS-89201, Tebubio, France), Sytox Green (S7020, Invitrogen, Belgium), Hoechst 33342 (H3570, Invitrogen, Belgium), LysoTracker Green DND-26 (L7526, Invitrogen, Belgium), MitoTracker Green (M7514, Invitrogen, Belgium), Propidium Iodide (25535-16-4, Sigma-Aldrich, Belgium), Cholera Toxin Subunit B (Recombinant) Alexa Fluor 488^TM^ Conjugate (C34775, Invitrogen, Belgium), Tetramethylrhodamine, methyl ester (TMRM) dye (T668, Invitrogen, Belgium). The following viability assay kits were used: CellTiter-Glo® Luminescent Cell Viability assay (G7571, Promega, Belgium), CellTiter 96® Aqueous Non-Radioactive Cell Proliferation Assay, (G5421, Promega, Belgium), and CellTiter^®^ 3D Cell Viability Assay (G9682, Promega, Belgium). Normalization of the ATP assays was done by Pierce BCA Protein assay kit (23227, Thermo Fisher Scientific Inc., Belgium). The following mitochondrial drugs were used: FCCP (Sigma-Aldrich C2920-10MG, Belgium), oligomycin A (75351, Sigma-Aldrich, Belgium), Rotenone (R8875-1G, Sigma-Aldrich, Belgium), and Antimycin A (A8674-25MG, Sigma-Aldrich, Belgium).

Spheroids were formed using the following methods: 0,5 wt% Lipidure^®^-CM5206^TM^ (AMS.52000034GB1G, Amsbio, UK) coating on U-bottom 96-well plate (10062-900, VWR, Belgium), BIOFLOAT^TM^ (83.3925.400, Sarstedt, Belgium), 3D Petri Dish^®^ micromolds (Z764000-6EA, MicroTissue Inc., USA), SphericalPlate 5D^®^ 24-well (SP5D-24W, Kugelmeiers, Switzerland) and micro-patterned 3D-printed PDMS stamps^63^ producing 1589 spheroids per mold (provided by the Centre for Microsystems Technology, Professor Dr. Jan Vanfleteren, Ghent University).

Other reagents included glucose oxidase (250 µg/ml), KH_2_PO_4_ (4873, Merck, Belgium), Na_2_SO_3_ (239321-500G, Sigma-Aldrich, Belgium), 69 kDa dextran (D-1537 Sigma-Aldrich, Belgium), HEPES-Na (25249, Serva Electrophoresis, Germany), NaCl (CL00.1429.100, Chemlab, Belgium), EDTA (E5134-100G, Sigma-Aldrich, Belgium), 1% Igepal (CA630, Sigma-Aldrich, Belgium), and protease inhibitor cocktail (D2714, Sigma-Aldrich, Belgium).

### Design and characterization of the nanoparticle probes

Oxygen indicator platinum(II) meso-tetra(4-fluorophenyl)tetrabenzoporphyrin (PtTPTBPF)^72^ was prepared *via* the template method reported by Hutter *et al.*^111^ The reference dye (BF2 chelate of (3,5-diphenyl-1H-pyrrol-2-yl)(3,5-diphenylpyrrol-2-ylidene)amine) “aza-BODIPY” was synthesized according to the published protocol.^74^ ^1^H NMR spectra for both dyes are provided in Supporting Information (Figures S1 and S2). The nanoparticles were prepared as described previously.^71^ Briefly, the dyes and the polymer were dissolved in organic solvent (acetone for RL-100 and acetone: tetrahydrofuran (9:1 v/v) for PMMA-MA, 0.2% wt. solution in both cases) and 5x volume of deionized water was added rapidly (1-2 seconds) under vigorous stirring. The organic solvents were removed under reduced pressure.

**Luminescence spectra** were recorded on a Fluorolog-3 luminescence spectrometer (Horiba, Germany) equipped with a NIR-sensitive R2658 photomultiplier (Hamamatsu, Germany). The measurements were performed in 10 mm screw-capped quartz cuvettes (Starna Scientific, UK). Oxygen concentration was adjusted by bubbling nitrogen (99.999% purity, Air Liquide, Austria) or its mixtures with compressed air through the dispersion of the particles. The gas mixtures were obtained using mass-flow controllers from Voegtlin (Switzerland, red-y smart series). The concentration of particles was adjusted to ∼50 mg/L to avoid the inner filter effect. To investigate the effect of serum on the luminescence, the dispersions containing 10% volume of serum were measured in the same cuvette (Starna Scientific, UK). To avoid foam formation oxygen concentrations were adjusted by introducing the gas mixtures for 1h into the head-space of the cuvette and stirring the solution with a magnetic stirring bar.

**Luminescence decay times** were measured on the same spectrometer in the time domain mode using a 455 nm SpectraLED (Horiba, Germany) as an excitation source (for phosphorescence of PtTPTBPF) or 635 nm NanoLED (Horiba, Germany) as an excitation source (for fluorescence of aza-BODIPY) and a DeltaHub module (Horiba, Germany). Data analysis was performed on DAS-6 Analysis software from Horiba.

#### Transmission electron microscopy

Morphology and size distribution were investigated for the three production batches of MMIR1 and MMIR-by drop casting (20 µg/ml, 2 µl) and dried overnight on Formvar/Carbon-coated hexagonal copper mesh grids (FCF200H-CU-TB, Electron Microscopy Sciences). The grids were observed on a transmission electron microscope JEM 1010 (Jeol, Ltd, Japan) equipped with a charge-coupled device side-mounted Veleta camera (Emsis, Germany). Nanoparticle size was measured manually using ImageJ software (NIH, USA).

#### Nanoparticle tracking analysis

Nanoparticle tracking analysis (NTA) was performed for the three batches of MMIR1 and MMIR-using a NanoSight LM10-HS microscope (NanoSight, Amesbury, UK) equipped with a 45 mW 488 nm laser and an automatic syringe pump system. Three 30s videos were recorded of each sample with camera levels of 13 (for MMIR-) and 15 (for MMIR1, a detection threshold of 3, and a syringe pump infusion speed of 20. Temperatures were monitored throughout the measurements; we assumed a medium viscosity of 0.929 cP and the videos were analyzed with NTA software 3.4. For optimal measurements, the samples were diluted with Milli-Q water until particle concentration was within an optimal concentration range of the NTA Software (3×10^8^ – 1×10^9^ particles/ml). All size distributions determined with NTA correspond to the hydrodynamic diameters of the particles in suspension.

### Cell culture

Human colon cancer HCT116, human pancreatic cancer PANC1, human embryonic kidney cells HEK-293T, and ovarian cancer SKOV3 were obtained from Laboratory of Experimental Cancer Research, Ghent University. Epithelial breast cancer MCF7 cell line was obtained from Radiobiology lab, Ghent University. HCT116 wild type and HCT116 SCO_2_^-/-^ cell lines^98^ were obtained from Prof. P. Hwang (Cardiovascular and Cancer Genetics, National Institutes of Health). Human dental pulp stem cells (hDPSCs, PT-5025) and human umbilical vein endothelial cells (HUVEC, C2519A) were from Lonza (Belgium). A short tandem repeat (STR) showed differences in both obtained HCT116 cell lines, compared to the ATCC one (Table S1). Therefore, HCT116 ODW (from LECR and ATCC) and HCT116 Hwang (Prof. P. Hwang) are treated as separate cell lines. All presented data were produced with HCT116 ODW, until otherwise mentioned. All cell lines were cultured in the recommended medium (Table S2) and under standard conditions *i.e.*, a humidified atmosphere of 37 °C, 18.6% O2, and 5% CO2. At 80– 90% confluency cells were passaged using 0.25% trypsin/EDTA (25300104, Gibco, USA) solution. Cells were cultured and analysed under antibiotic-free conditions. Cells were regularly tested for lack of mycoplasma contamination using PCR (Eurofins Genomics).

#### Spheroid formation methods

HCT116 spheroid (initial 500 cells per spheroid) were seeded together with or without (for unstained samples) 10 µg/ml MMIR1 on the low-attachment methods: 0,5 wt% Lipidure-CM5206^TM^ coating on U-bottom 96-well plate, or BIOFLOAT^TM^ and the high throughput-methods 3D Petri Dish® micromolds producing 81 spheroids per mold, SphericalPlate 5D^®^ 24-well producing 750 spheroids per mold and micro-patterned 3D-printed PDMS stamps^63^ producing 1589 spheroids per mold. Spheroid formation was performed over 5-6 days, before transferring on a prepared microscopy dish, counter-staining, and microscopy.

#### Evaluation of MMIR nanoparticles with monolayer cell cultures

2D cultures were seeded on microscopy dishes at a density of 12,500 ∼ 50,000 cells/ well and incubated overnight for attachment. Loading of 2D cells with the oxygen sensing probes was typically performed by probe addition of 5-20 µg/ml incubation (17 h) in triplicate in a 5% CO2 incubator at 37°C, followed up by washing (3x) with growth medium, prior to microscopy and other experiments. **For intracellular localization,** HCT116 were stained with MMIR probes (5-20 µg/ml, overnight), followed up by the 1 h co-staining either with 25 ng/ml Calcein Green-AM, 100 nM MitoTracker Green FM, 200 nM LysoTracker Green DND-26, 2 nM HXT33342 or 2 µM Cholera Toxin Subunit B (Recombinant) Alexa Fluor 488^TM^ Conjugate, washing and imaging on a confocal microscope.

### Assessment of cytotoxicity

The cytotoxicity effects of various concentrations of nanoparticles for 2D cultures were investigated by co-staining with 30 nM Sytox green and 0.5 µM Hoechst 33342 (1 h), washing and imaging on a widefield fluorescence inverted microscope IX81 (Olympus). Viability effects on the membrane integrity were calculated by dividing N_viable cells_ by total cell number (viable + dead cells).**Effects on total cell ATP (resting and metabolic stimulation)** were probed using CellTiter-Glo® Luminescent Cell Viability assay (Promega), according to the manufacturer’s protocol with the luminescence recorded using the Spark multimode microplate reader (Tecan Spark 20M). Luminescence data on ATP were normalized by extracting the total cell protein with PEB buffer (50 mM HEPES-Na, pH 7.4, 150 mM NaCl, 1 mM EDTA, 1% Igepal CA630, and protease inhibitor cocktail and analysed with Pierce BCA Protein assay kit as described previously^112^.

Viability of 2D cell cultures was also studied using colorimetric 3-(4,5-dimethylthiazol-2-yl)-5-(3-carboxymethoxyphenyl)-2-(4-sulfophenyl)-2H-tetrazolium **(MTS) assay** (CellTiter 96® Aqueous Non-Radioactive Cell Proliferation Assay, according to the manufacturer’s protocol, with the absorbance at 490 nm read using Universal Microplate Reader EL800 (Bio-TEK Instruments, Inc., USA). Linearity of response was investigated by seeding cells over the range of 64,000∼2,000 cells/well. For all 2D viability studies, 30,000 cells/well were seeded.

Cytotoxicity effects on MMIR-stained spheroids were investigated using the CellTiter^®^ **3D Cell Viability** Assay. Briefly, HCT116 spheroids were formed on 3D Petri Dish® micromolds, at a density of 500 cells/ well and grown for 5 days. During formation, 20 µg/ml MMIR1 was added to 10 replicates. Once formed, spheroids were transferred into a 96-well flat bottom plate coated with 0.07 mg/ml collagen IV/ 0.015 mg D-poly-lysine and allowed to attach for 2 hours. CellTiter-Glo 3D reagent was added in equal amounts of cell culture medium and lysis by shaking for 5 minutes. After stabilizing the luminescent signal for 25 minutes, the signal was read using the Spark multimode microplate reader (Tecan Spark 20M) with an integration time of 1 second per well. Results were normalized by spheroid size (area square) division. Unstained and stained spheroids were also tested for differences in their NAD(P)H using two-photon microscopy (Fig. S7).

### Fluorescence microscopy

#### Widefield fluorescence microscopy

Fluorescence microscopy was performed on a widefield fluorescence inverted microscope IX81 (Olympus), with motorized Z-axis control, CoolLED pE4000 (16 channels, 365-770 nm), ORCA-Flash4.0LT+ (Hamamatsu) cMOS camera, glass warming plate Okolab, CellSens Dimension v.3 software and air objectives 40x/0.6 UPlanFL N and 20x/0.45 UPLanFL N. The CellSens Dimension (Olympus) software was used for imaging acquisition and processing with fixed settings (Table 1).

**Table 1:** Image acquisition widefield fluorescence inverted microscope (Olympus)

| Dye | Light source | Excitation filter (nm) | Emission filter (nm) | Exposure time (ms) | Power (%) | Objective |
| --- | --- | --- | --- | --- | --- | --- |
| HXT 33342 | 405 nm | 391-437 | 420-460 | 2 | 50 | UPlanFL N 40x/0.6 air |
| SYTOX Green | 470 nm | 460-495 | 510-550 | 10 | 50 | UPlanFL N 40x/0.6 air |
| MMIR1 (reference) | 580 nm | 545-580 | 610 nm longpass | 20 | 25 | UPlanFL N 40x/0.6 air |
| MMIR1 (O<sub>2</sub> sensitive) | 635 nm | None/ 685 nm longpass | 705-845 nm | 20 | 25 | UPlanFL N 40x/0.6 air |
| Calcein Green-AM | 490 nm | 460-495 | 510-550 | 20 | 50 | UPlanFL N 40x/0.6 air |
| Propidium Iodide | 550 nm | 545-580 | 610 nm longpass | 25 | 25 | UPlanFL N 40x/0.6 air |
| TMRM | 550 nm | 545-580 | 610 nm longpass | 5 | 25 | UPlanFL N 20x/0.45 |
| NADPH<sup>113</sup> | 460 nm | 460-495 | 510-550 | 120 | 25 | UPlanFL N<br>40x/0.6 air |
| FAD | 460 nm | 460-495 | 510-550 | 40 | 50 | UPlanFL N<br>40x/0.6 air |

**Photostability** properties of the nanoparticles NPs were investigated on live HCT116 spheroids using the widefield fluorescence microscope IX81 (Olympus). The photostability experiment was done by calculating the percentage of the reference intensity after 10 cycles of illumination for 10 spheroids. 10 nM of orange-red mitochondrial TMRM dye was used as a reference. **Kinetic responses** to mitochondrial stimuli on 5 days old HCT116 spheroids (formed Microtissue^®^ molds, 500 cells per spheroid) were investigated by sequential treatment with mitochondrial uncoupler FCCP (1 µM) and mitochondrial inhibitor Rotenone (1 µM). The ratio responses were further validated by oxygenating HCT116 spheroids with rotenone (1 µM) and Antimycin A (1 µM), followed up by deoxygenation using glucose oxidase (250 µg/ml) and potassium sulfite solution (PSS) containing 50 mg/ml KH_2_PO_4_ and 50 mg/ml Na_2_SO_3_.

**To visualize oxygen gradients**, ratiometric analysis the following formula was used: R = 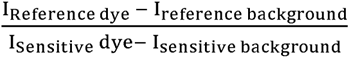 for each pixel of the 3D model. Each “ratio pixel” was converted into a colour gradient. We used the parameters “oxygen gradient” (R_periphery_ - R_core_) and “steepness” (ΔRatio/spheroids radius) introduced previously^63^.

#### Size dependent oxygenation

HCT116 cells were prestained with 10 µg/ml MMIR1 probe overnight before spheroid formation on 0,5 wt% Lipidure^®^ 96-well plates with addition of 1 µg/ml MMIR1 in multiple seeding densities (500, 1,000, 5,000 and 10,000 cells per well) in 6 replicates. Line profiles were taken using the “line profile” tool with CellSens software (Olympus).

#### Effect of medium on oxygenation

To investigate the effects of increased viscosity and glucose concentration on the oxygen gradient in live HCT116 spheroids, culture media was exchanged with imaging media with either D-glucose (0 – 25 mM) or 0 to 5%w/w 69 kDa dextran resulting in a viscosity ranging from 0.7748 cP to 2.25 cP at 37°C, 4 hours before imaging^114^. HCT116 spheroids were formed using the high throughput agarose micromolds method with the addition of MMIR1 (10 µg/ml) and grown for 5 days.

#### Inverted oxygenation gradient analysis

Live DPSCs spheroids were formed by seeding 300,000 and 1.300,000 cells/ml on a micro-patterned agarose-coated tissue culture plate^63^. Theoretically, the cells were then equally distributed among the 1585 microwells, resulting in 189 and 820 cells per spheroid, respectively. Live HCT116 spheroids were formed by seeding 500, 1,000, 10,000, and 20,000 cell per spheroid in a Lipidure^®^-coated plate (Amsbio). After 7 days (HCT116) and 8 days (DPSCs) of CO_2_ incubation at 37°C, spheroids were transferred to pre-coated microscopy dishes. Multi-parametric analysis was performed by PI staining (1 µg/ml, 1 h), NAD(P)H-specific fluorescent probe^113^ (20 µM, 1 h), and FAD autofluorescence (exc. 460 nm, 510-550 nm emission) on the Olympus IX81 microscope.

**Confocal FLIM microscopy** was performed on an inverted Stellaris 8 Falcon (Leica) microscope (Ghent Light Microscopy Core), equipped with the white-light laser (440-790 nm), HC PL Apo 10x/0.4 air, HC Fluotar 25x/0.95 W, HC PL Apo 40x/1.25 GLYC corr., HC PL Apo 63x/1.4 Oil objectives, HyD X, HyD R and HyD S detectors, temperature-controlled incubator, and dedicated LAS X acquisition and analysis software (ver. 4.6.0), as described previously^115^. For co-localisation studies, typically 40x/1.25 GLYC corr. objective was used, with MMIR excited at 614 nm (4.34%), emission collected at 639-696 nm (reference channel, HyD X3 detector) and 724-780 nm (O_2_-sensitive channel, HyD R detector), pixel dwell time 4.1 μs, 1024×1024 resolution. Counter-stains were imaged accordingly to their spectral settings.

**Two-photon FLIM microscopy** was performed on an inverted SP8 Falcon (Leica) microscope, equipped with IR Mai Tai HP laser (690-1040 nm), a temperature-controlled incubator, HC Fluotar 25x/0.95 W objective and dedicated LAS X acquisition and analysis software. NAD(P)H was excited at 740 nm (20% laser power) with emission collected at 420-470 nm, at 512×512 resolution, pixel dwell time 3.2 μs. Produced images were analysed using LAS X FLIM analysis suite for reconstructing 3D FLIM and phasor FLIM images. NAD(P)H-FLIM-based analysis of OxPhos / Glycolysis states in spheroids was done by comparing the shift of the NAD(P)H phasor plots of spheroids from the theoretical position of monoexponential free-NAD(P)H lifetime (∼0.45 ns)^116^ on the universal circle. Threshold 5, wavelet filter were applied for phasor plots corresponding to individual spheroid ROIs, and the theoretical position of free-NAD(P)H lifetime on a universal circle was determined in LAS X-FLIM software. Phasor plots were exported as TIFF files (600×1024 pixels) together with the circular mask and free-NAD(P)H position. Narrowed phasor plot clouds and free-NAD(P)H position on a universal circle were selected in ImageJ^117^ by application of corresponding threshold filters with fixed colour space, brightness, and saturation parameters to generate corresponding ROI masks. The centroids of phasor plot masks were determined using standard ‘measure’ protocol and their coordinates (in pixels), corresponding to their coordinates inside the universal circle were exported in Excel table format. Distance (*d*) calculation between centroids and theoretical free-NAD(P)H position was performed using Pythagorean formula

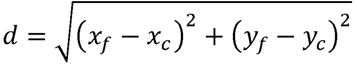

Where x_f_ and y_f_ –coordinates of free-NAD(P)H and x_c_ and y_c_ – coordinates of phasor plot centroids. The linear alignment of centroids with free-NADPH was verified by linear fitting with R^2^ = 0.96.

### Data assessment and statistics

2D cell staining studies were analysed by averaging the intensity values of 10 randomly selected ROI per replicate. For spheroids, 10 ROIs at the periphery and 10 ROIs at the core were randomly selected. All MMIR application methods were performed in at least 3 independent experimental replicates to ensure reproducibility. All results are represented as mean values ± SEM. The number of replicates can be found in the legend of each figure. Figures were made using Inkscape software. Quantitative data were analysed, and graphs were made using GraphPad Prism 9. Normally distributed data were subjected to a Student’s *t*-test or a one-way ANOVA. Alternatively, the non-parametric Mann-Whitney U test (two groups) or Kruskal-Wallis test (multiple groups) were performed. In case of a significant result, the Tukey HSD test (parametric) or a pairwise comparison through the Wilcoxon rank sum test with Bonferroni correction was performed. Significant differences are shown as followed: * = P < 0.05, ** = P<0.01, *** = P<0.001 and **** = P<0.0001.

## Supporting information

Supplementary Figures and tables combined

## Acknowledgements

We would like to thank O. De Wever and A. Baeyens (Ghent University) for sharing the cell lines, P. Hwang (National Institutes of Health) for HCT116 wild type and KO SCO_2_^-/-^ cell lines, M. Claeys (The Nematology Research Unit, UGent TEM Core Facility) for help with transmission the electron microscopy, D. Krysko (Ghent University) and A. Labro for support with a multimode microplate reader, C. Scheele (VIB, KU Leuven) for support with multiphoton microscopy and M. Barroso lab (Albany Medical College) for discussions on 3D models and O_2_ probes.

This research was supported by the Special Research Fund (BOF) starting investigator grant (BOF/STA/202009/003), Research Foundation Flanders (FWO) grants (I001922N, K1D0222N, K1DKE23N, K207223N, K1DKX23N, K1D6622N), European Union, fliMAGIN3D-DN Horizon Europe MSCA-DN No.101073507 and the Austrian Science Fund FWF, grant no. P32079-N37.

## Supporting Information

### Supporting information available

**Supplementary figure S1:** ^1^H NMR spectrum (CDCl_3_, 400 MHz) of BF_2_ chelate of (3,5-diphenyl-1H-pyrrol-2-yl)(3,5-diphenylpyrrol-2-ylidene)amine (“aza-BODIPY”).

**Supplementary figure S2:** ^1^H NMR spectrum (CD_2_Cl_2_, 400 MHz) of PtTPTBPF.

**Supplementary figure S3:** Normalized absorption spectra of the particles (1:1 wt. ratio of the dyes) for 3 different batches of each type.

**Supplementary figure S4:** Emission spectra (λ_exc_ = 615 nm) of the particles (1:1 wt. ratio of the dyes) for three different batches of each type.

**Supplementary figure S5:** Appearance of aqueous dispersions of the nanoparticles (concentration 2 mg/mL). From left to right: MMIR after two weeks storage at 4 °C, MMIR1 after

∼2 years storage at 4 °C, MMIR-after 2 weeks storage at 4 °C, MMIR-after ∼2 years storage at 4 °C.

**Supplementary figure S6:** Emission spectra of MMIR1 and MMIR-nanoparticles (dye ratio 1:1) in anoxic water. The absorption of both dispersions was identical at the excitation wavelength (615 nm).

**Supplementary figure S7:** Luminescence decays of the reference dye in MMIR1 and MMIR-beads (both 1:1 ratio of the dyes) monitored at 660 nm (λ_exc_ = 635 nm, NanoLED, Horiba; 23 °C).

**Supplementary figure S8:** Phosphorescence decay of PtTPTBPF in MMIR1 beads (1:1 ratio of the dyes) monitored at 760 nm (λ_exc_ = 456 nm, SpectraLED from Horiba; 23 °C).

**Supplementary figure S9:** Phosphorescence decay of PtTPTBPF in MMIR-beads (1:1 ratio of the dyes) monitored at 760 nm (λ_exc_ = 455 nm, SpectraLED from Horiba; 23 °C).

**Supplementary figure S10:** Nanoparticle size measurements of three production batches of MMIR1 and MMIR-using NanoSight (A) and TEM (B) methods.

**Supplementary figure S11:** Comparison of positively charged MMIR1 with different sensitive/reference dye ratios and negatively charged (1:1) MMIR-probe.

**Supplementary figure S12:** Fluorescence signals of MMIR1-stained HCT116 spheroids over 26 days period.

**Supplementary figure S13**: Intracellular localization of MMIR1 after overnight incubation with HCT116 cells (5 µg/ml, 17 h) shows endo- and lysosomal localization.

**Supplementary figure S14:** Cell line-dependent uptake of the positive (RL100) and negative (PMMA) charged MMIR probes.

**Supplementary figure S15:** Pre-staining of HCT116 cells overnight ensures uniform staining.

**Supplementary figure S16:** Addition of MMIR probes shows no significant cell death in monolayer and spheroids of HCT116 cells.

**Supplementary figure S17**: Viability assay using the CellTiter 96 Aqueous Non-Radioactive Cell Proliferation Assay (MTS, Promega) shows no cellular toxicity due to 24 h incubation of both cationic (MMIR1) and negatively charged (MMIR-) nanosensors with live HCT116 cells.

**Supplementary figure S18:** Two-photon FLIM of NAD(P)H shows no changes in cell redox (NAD(P)H fluorescence lifetimes) in response to MMIR1 addition to live HCT116 spheroids.

**Supplementary figure S19:** Dynamic response of MMIR probes to drugs affecting cell bioenergetics and deoxygenation.

**Supplementary figure S20:** Size-dependent oxygenation of live HCT116 cells spheroids.

**Supplementary figure S21:** Increased cell medium viscosity results in the formation of a more hypoxic core.

**Supplementary figure S22**: Decreased cell media glucose concentration does not affect spheroids oxygenation.

**Supplementary figure S23:** Formation methods affect the viability and oxygenation in live HCT116 spheroids.

**Supplementary figure S24**: Inverted gradient formation in live HCT116 spheroids is both time and size-dependent.

**Supplementary figure S25:** Analysis of “inverted” oxygenation gradient in live DPSCs and HCT116 spheroids.

**Supplementary figure S26:** Cell death comparison between “direct” and “inverted” gradients in DPSCs and HCT116 spheroids formed on respectively microagarose molds and Lipidure®-coated plates.

**Supplementary figure S27:** Examples of two-photon NAD(P)H-FLIM microscopy and phasor plots of HCT116 spheroids with different sizes.

**Table ST1**. TR authentication HCT116 cell lines

**Table ST2**. Composition of used growth and imaging media.

**Table ST3.** Morphological characterization of spheroid populations

