## Supplementary Figures and tables combined for "Live microscopy of multicellular spheroids with the multi-modal near-infrared nanoparticles reveals differences in oxygenation gradients"

**Contents**

Supplementary figures **S1-S27.**

Supplementary tables **ST1-ST3**.

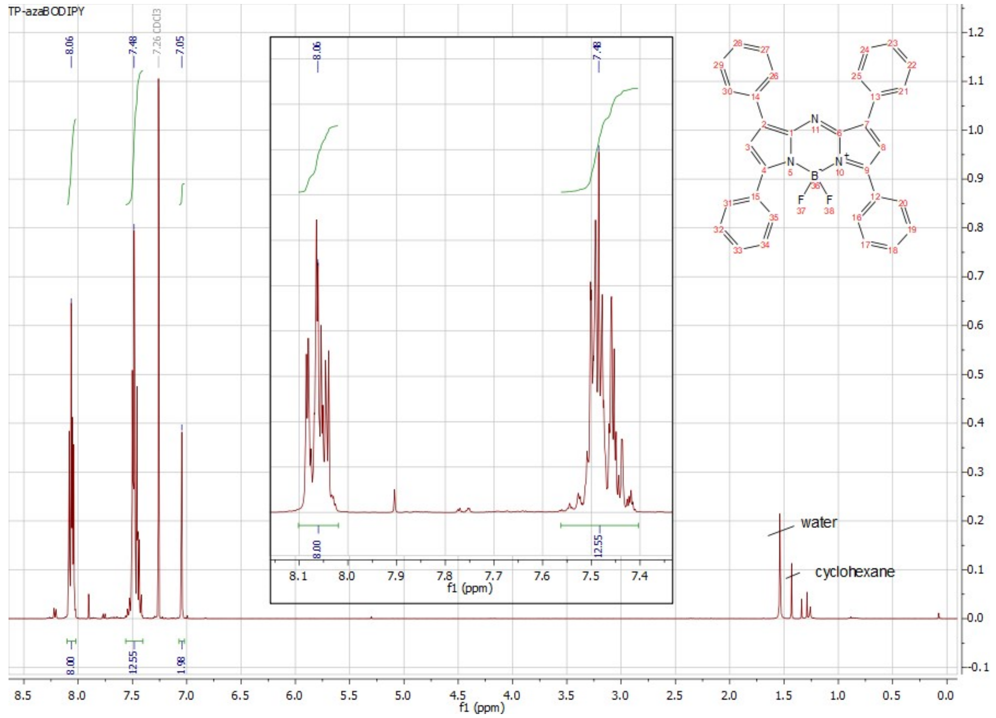


**Supplementary figure S1:** $^1$H NMR spectrum ($CDCl_3$, 400 MHz) of $BF_2$ chelate of (3,5-diphenyl-1H-pyrrol-2-yl)(3,5-diphenylpyrrol-2-ylidene)amine ("aza-BODIPY").

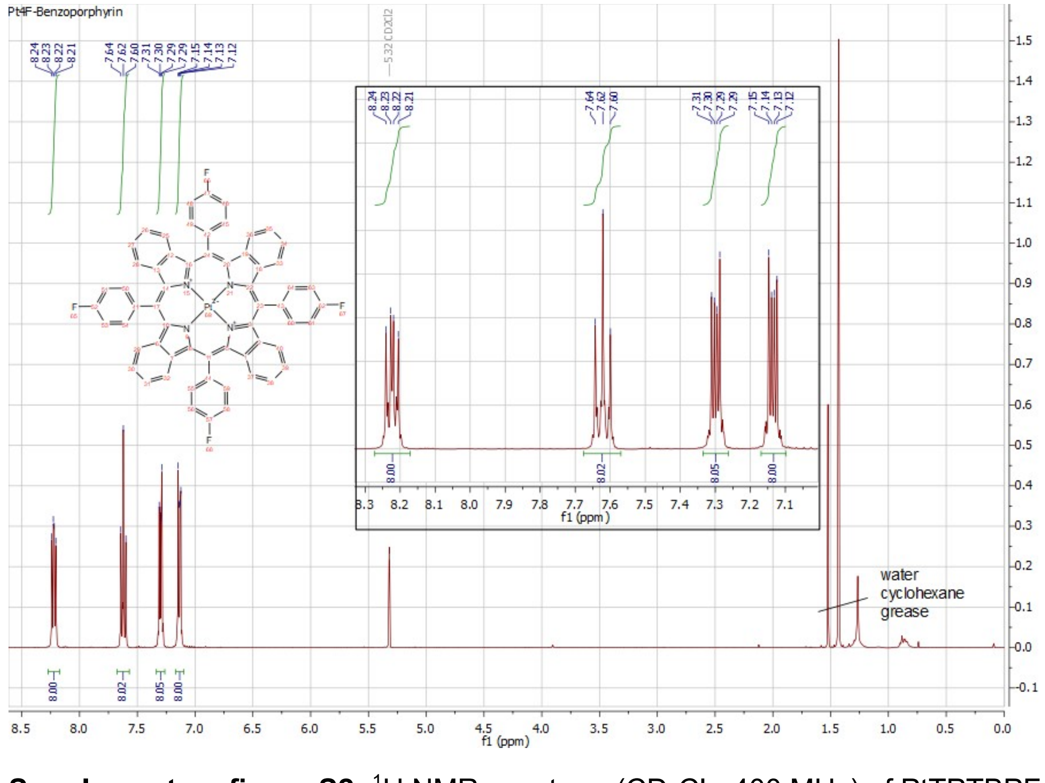


**Supplementary figure S2:** $^1$H NMR spectrum ($CD_2Cl_2$, 400 MHz) of PtTPTBPF.

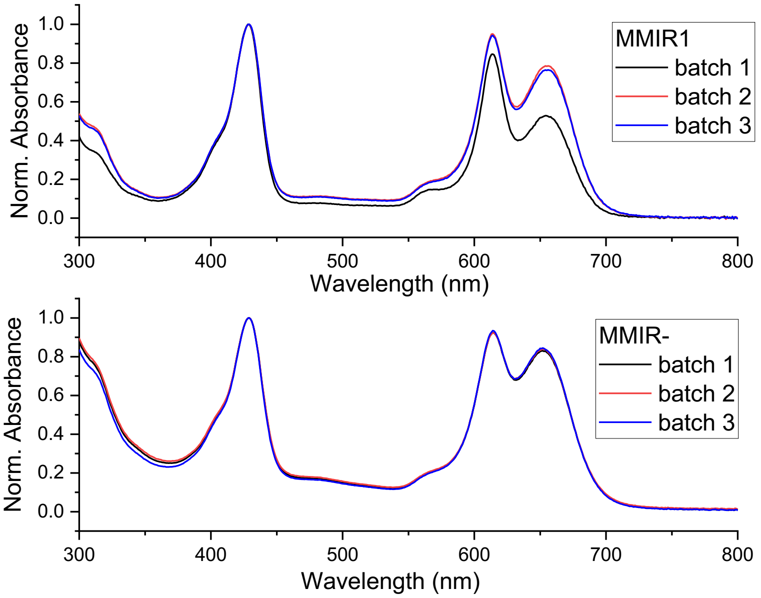


**Supplementary figure S3:** normalized absorption spectra of the particles (1:1 wt. ratio of the dyes) for 3 different batches of each type.

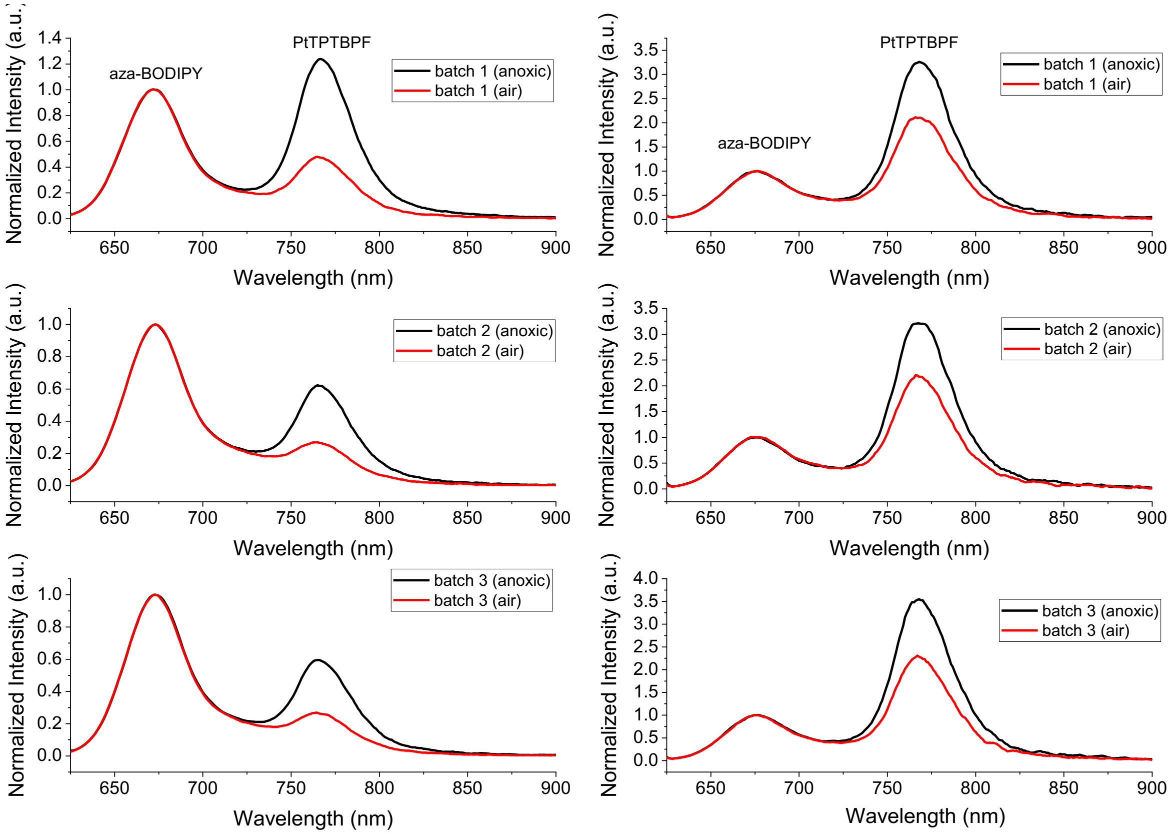


**Supplementary figure S4:** Emission spectra ($\lambda_{exc}$ = 615 nm) of the particles (1:1 wt. ratio of the dyes) for three different batches of each type. Left row: MMIR1, right row: MMIR- beads.

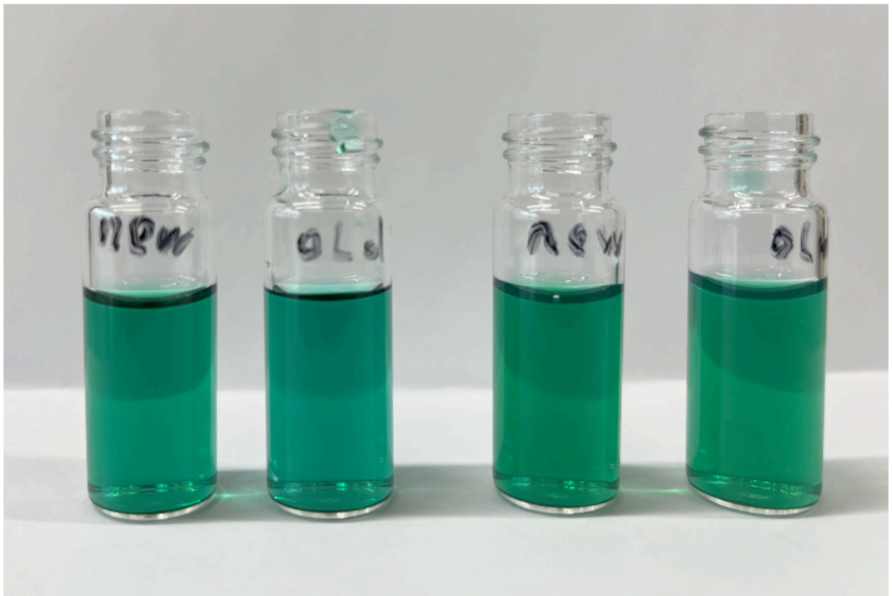


**Supplementary figure S5:** Appearance of aqueous dispersions of the nanoparticles (concentration 2 mg/mL). From left to right: MMIR after two weeks storage at 4 °C, MMIR1 after ~2 years storage at 4 °C, MMIR- after 2 weeks storage at 4 °C, MMIR- after ~2 years storage at 4 °C.

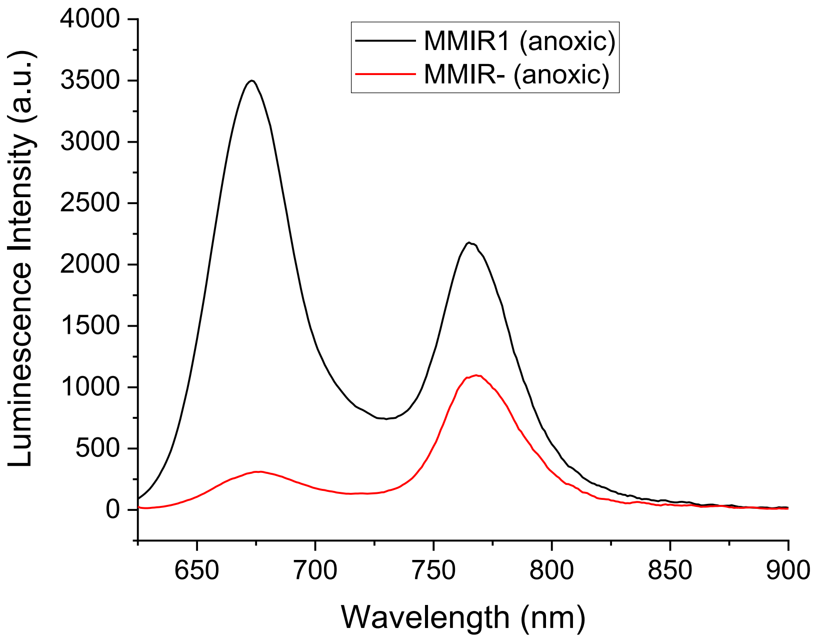


**Supplementary figure S6:** Emission spectra of MMIR1 and MMIR- nanoparticles (dye ratio 1:1) in anoxic water. The absorption of both dispersions was identical at the excitation wavelength (615 nm).

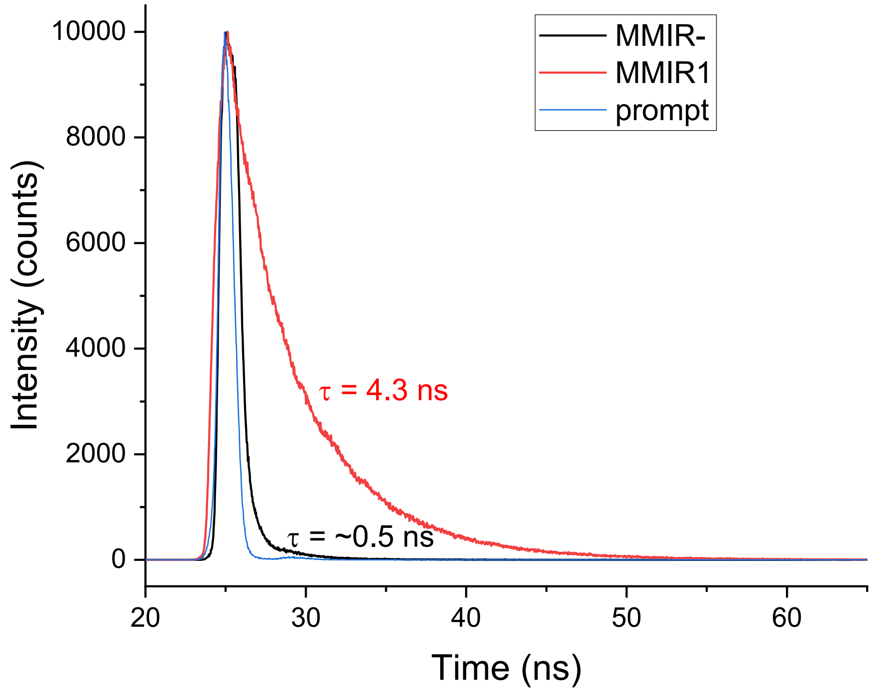


**Supplementary figure S7:** Luminescence decays of the reference dye in MMIR1 and MMIR- beads (both 1:1 ratio of the dyes) monitored at 660 nm ($\lambda_{exc}$ = 635 nm, NanoLED, Horiba; 23 °C).

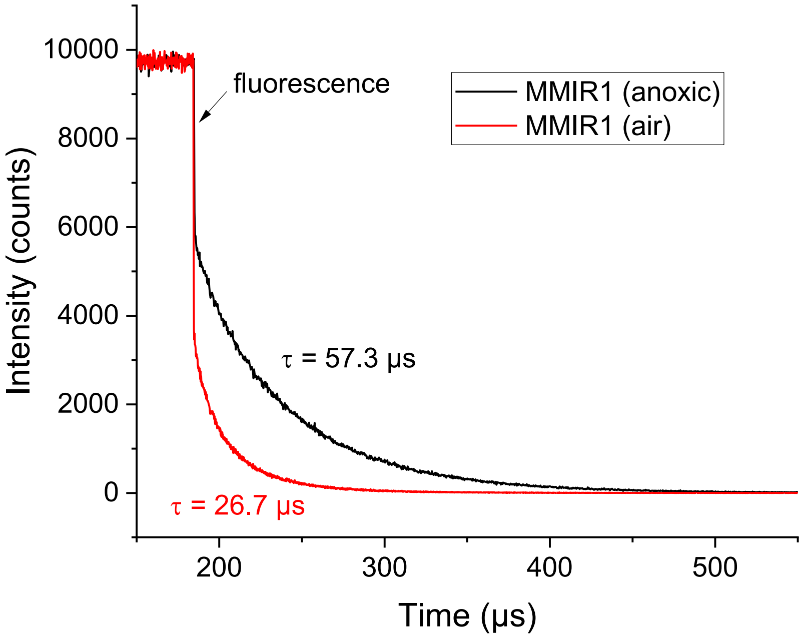


**Supplementary figure S8:** Phosphorescence decay of PtTPTBPF in MMIR1 beads (1:1 ratio of the dyes) monitored at 760 nm ($\lambda_{exc}$ = 456 nm, SpectraLED from Horiba; 23 °C).

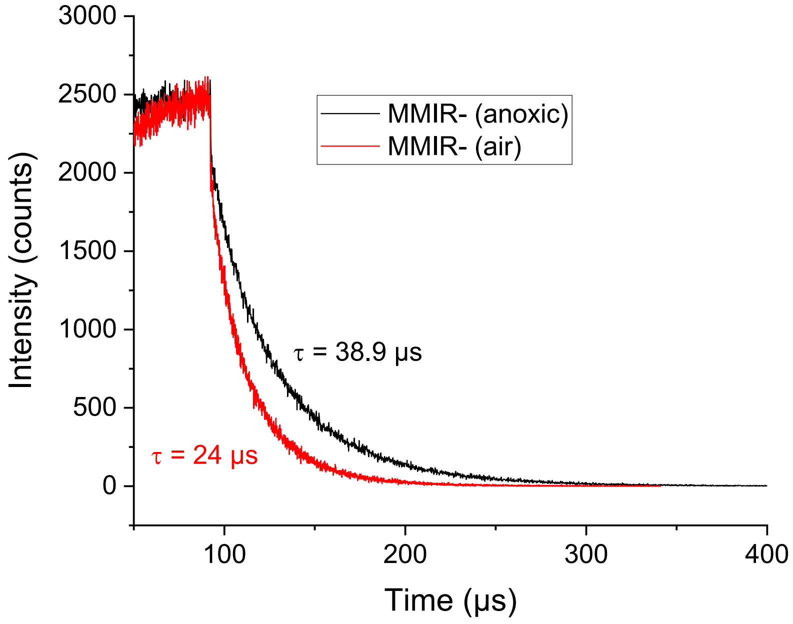


**Supplementary figure S9:** Phosphorescence decay of PtTPTBPF in MMIR- beads (1:1 ratio of the dyes) monitored at 760 nm ($\lambda_{exc}$ = 455 nm, SpectraLED from Horiba; 23 °C).

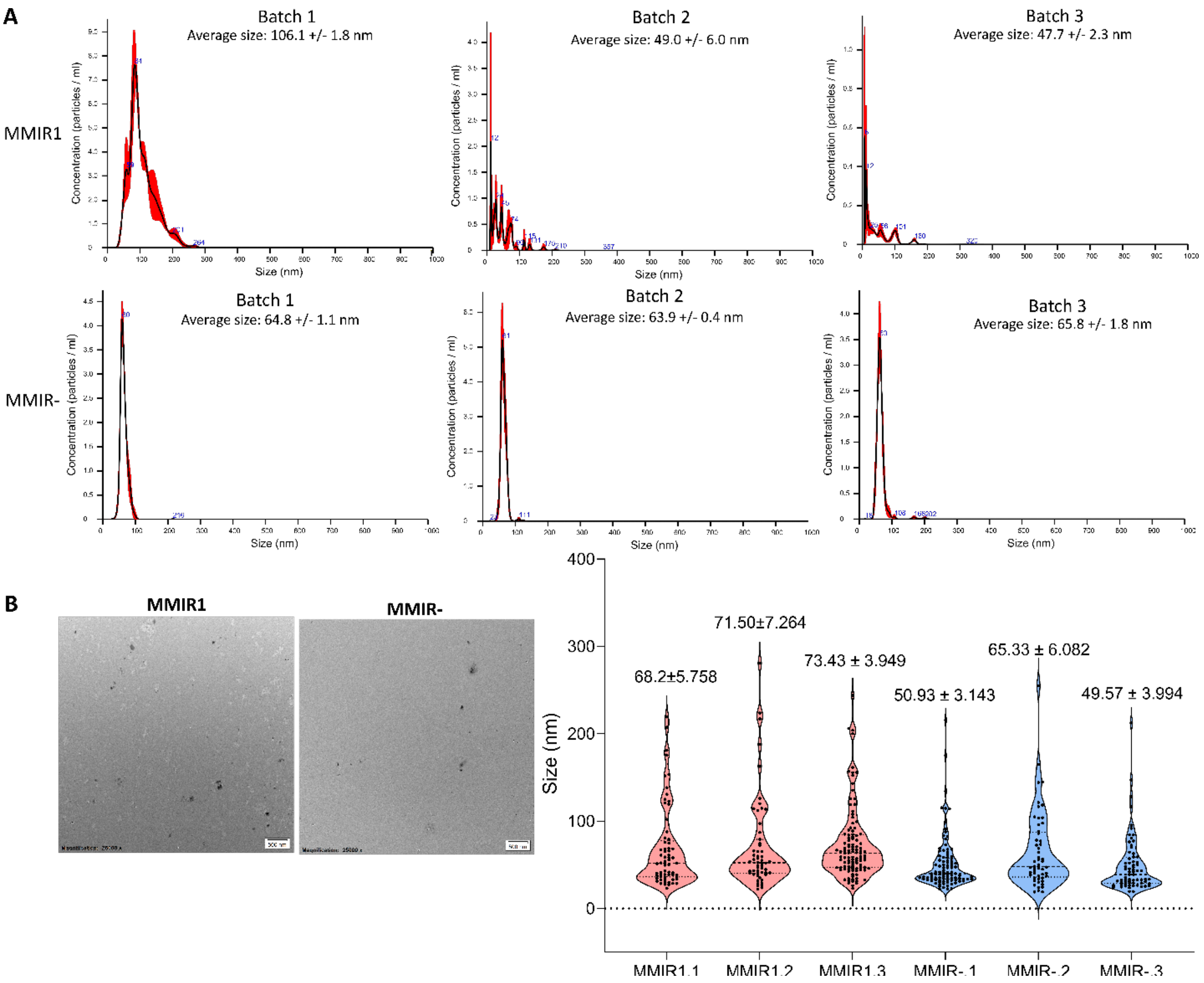


**Supplementary figure S10:** Nanoparticle size measurements of three production batches of MMIR1 and MMIR- using NanoSight (A) and TEM (B) methods. A: for each batch, three 30 second videos were used for the NanoParticle Tracking Analysis (NTA). Red error bars indicate ± SEM. B: Data shows for each batch the average size ± standard error of 53-110 counted nanoparticles. Scale bar is 200 nm.

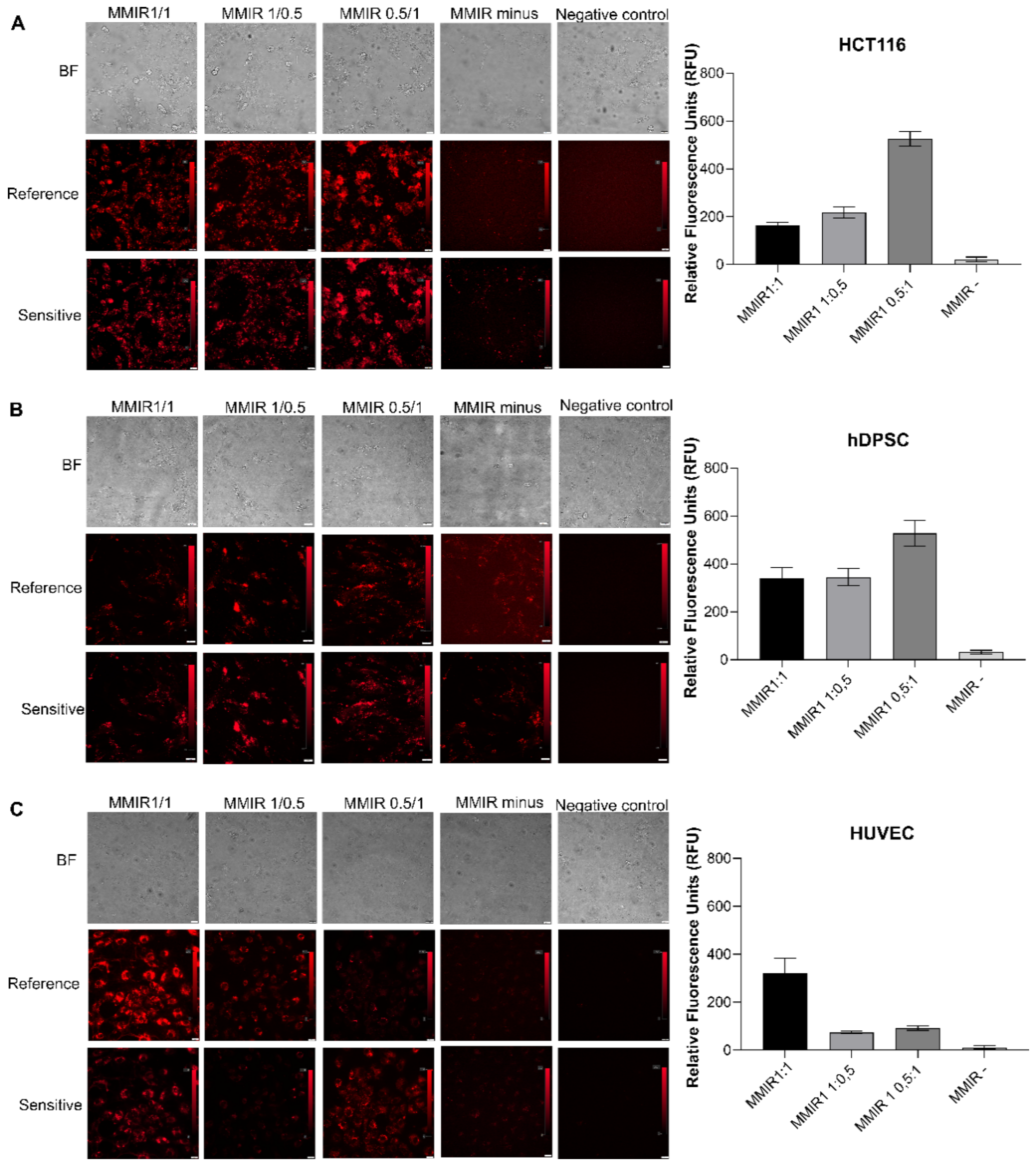


**Supplementary figure S11: Comparison of positively charged MMIR1 with different sensitive/reference dye ratios and negatively charged (1:1) MMIR- probe**. A: Fluorescence images of HCT116 cells stained overnight (5 µg/ml, overnight staining). B: Fluorescence images of human DPSCs cells stained overnight (5 µg/ml, overnight staining). C: Fluorescence images of HUVEC cells stained overnight (5 µg/ml, overnight staining). Results show brightfield (BF) images and both reference and sensitive fluorescence channels with scale bar (20 µm) and intensity bar. Relative fluorescence units (RFU) of the reference dye is shown for each cell line on the right. Results show the average ± standard error of 3 replicates.

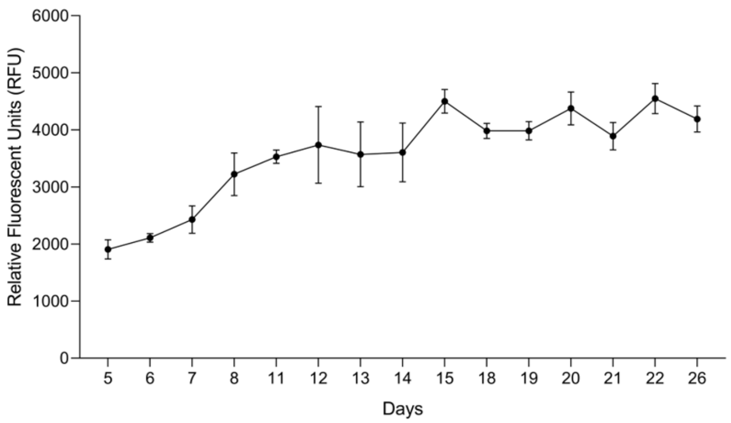


**Supplementary figure S12:-Fluorescence signals of MMIR1-stained HCT116 spheroids over 26 days period.** Formation on an ultra-low attachment 96-well plate with addition of MMIR (10 µg/ml) and imaged using widefield fluorescence inverted microscope IX81 (Olympus). Data show the mean ± standard error for 3 spheroids (reference channel).

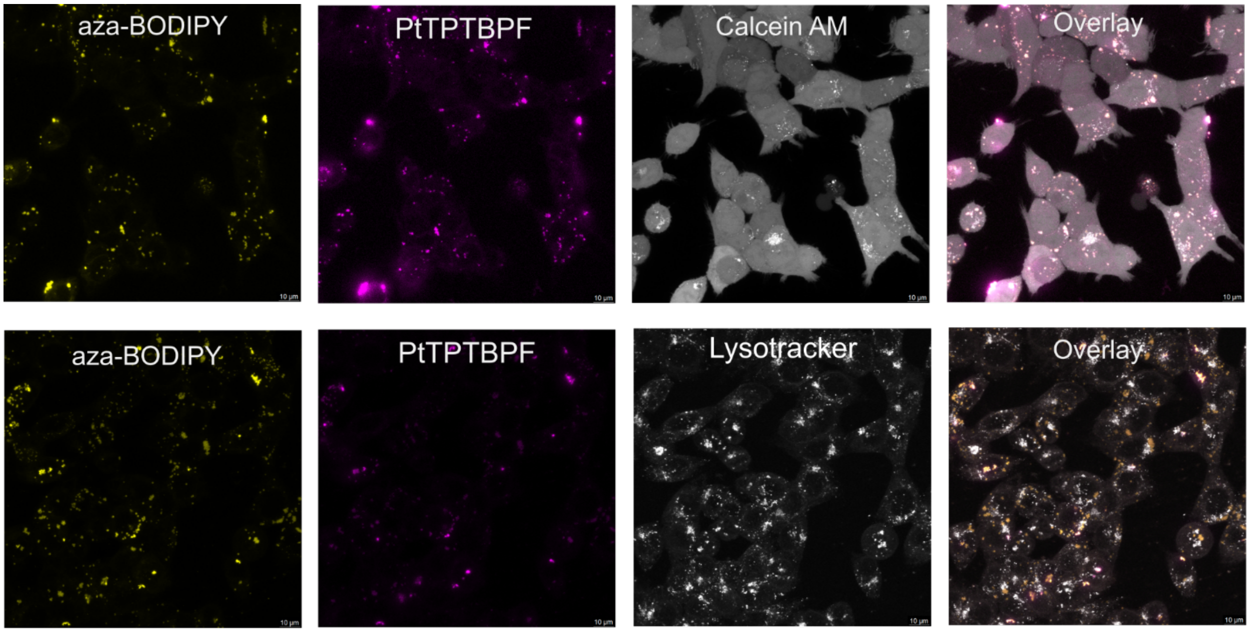


**Supplementary figure S13: Intracellular localization of MMIR1 after overnight incubation with HCT116 cells (5 µg/ml, 17 h) shows endo- and lysosomal localization.** Cells were co-stained for 2 hours with Calcein Green (25 ng/ml) and LysoTracker Green (200 nM). Confocal single optical sections are shown. Scale bar is 10 µm.

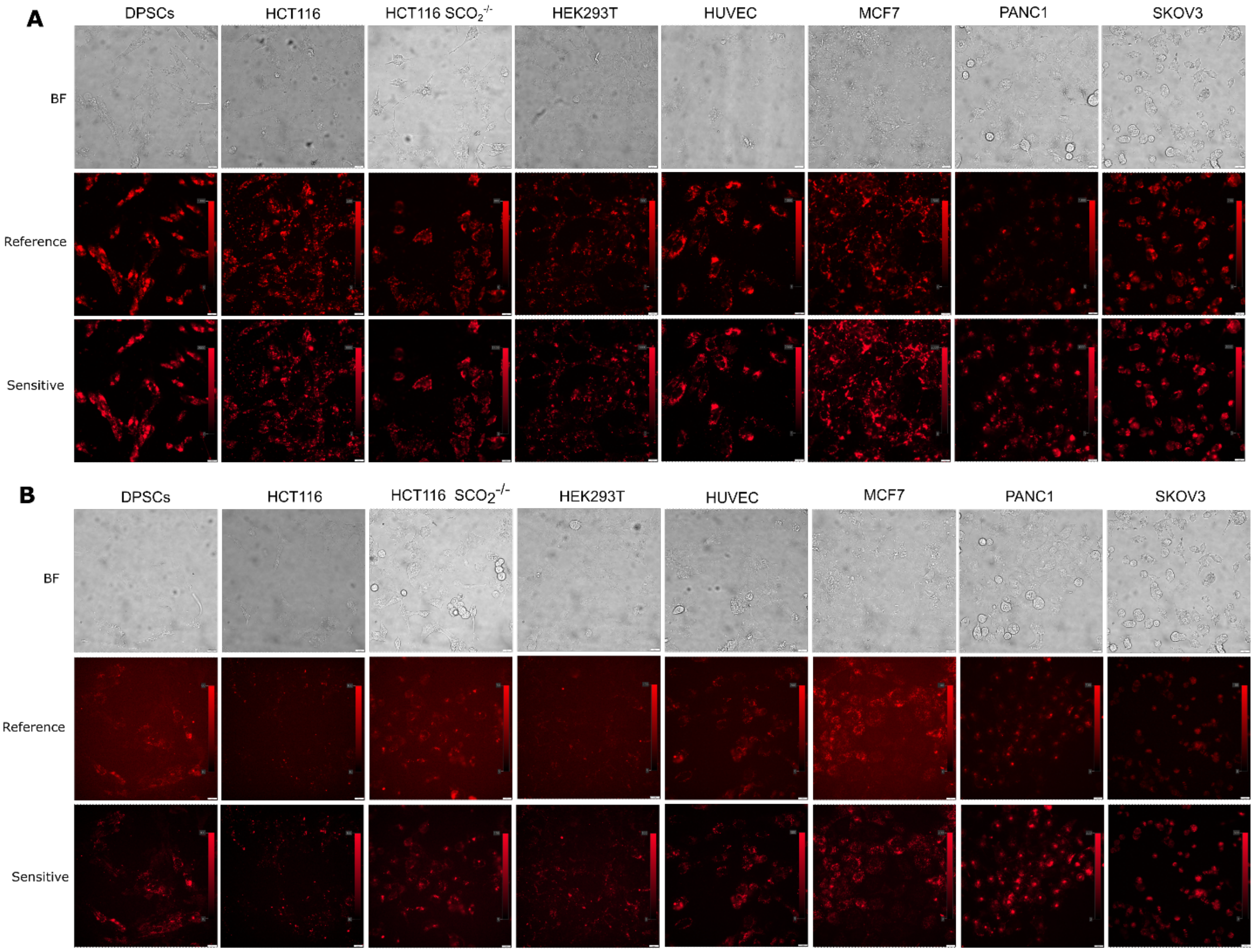


**Supplementary figure S14: Cell line-dependent uptake of the positive (RL100) and negative (PMMA) charged MMIR probes.** A: Fluorescence images of MMIR (5 µg/ml). B: Fluorescence images of MMIR- (5 µg/ml). Results show brightfield (BF) images and reference and sensitive fluorescence channels with scale bar (20 µm) and intensity bar. C: The relative fluorescence units of the reference dye in the RL100 NP and PMMA NP. Data shown is an average of 3 repeats (with background subtraction) ± SEM. DPSCs: dental pulp stem cells, HCT116: human colon cancer cell line, HEK293T: Human embryonic kidney cells, HUVEC: human umbilical vein endothelial cells, MCF7: epithelial metastatic adenocarcinoma, PANC1: pancreas epithelioid carcinoma, SKOV3: ovarian adenocarcinoma and RFU: Relative fluorescence units

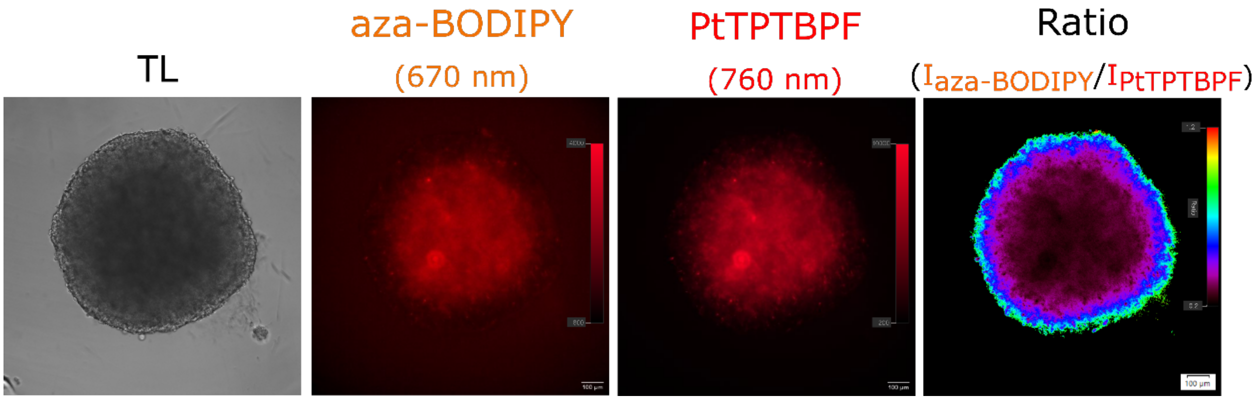


**Supplementary figure S15: Pre-staining of HCT116 cells overnight ensures uniform staining.** HCT116 cells were stained overnight with MMIR1 (50 µg/ml, 17 h), before formation using Lipidure®-coated plates. Imaged using widefield fluorescence inverted microscope IX81 (Olympus). Scale bar is 100 µm.

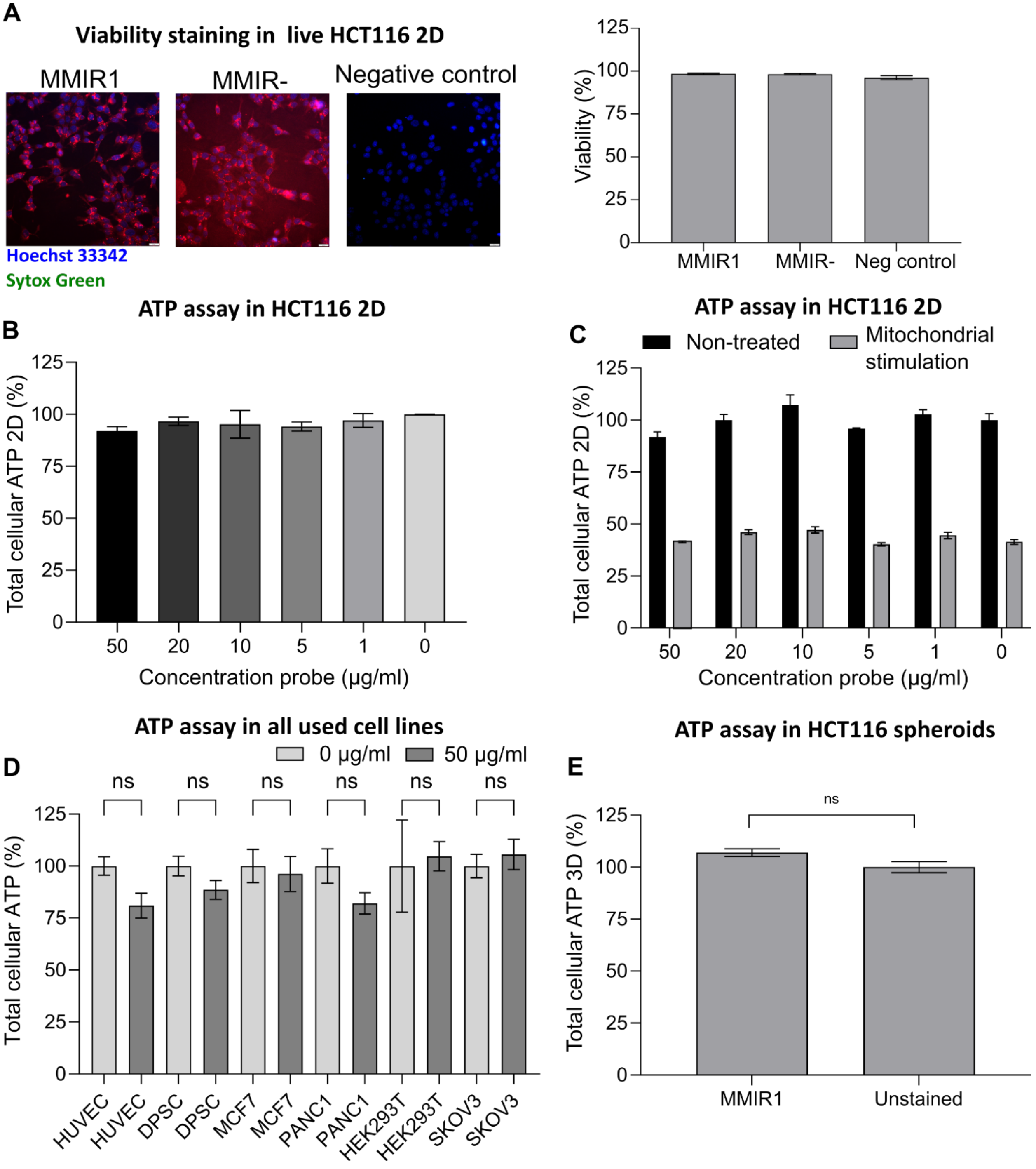


**Supplementary figure S16: Addition of MMIR probes shows no significant cell death in monolayer and spheroids of HCT116 cells.** A: Merged fluorescent images of HCT116 cells stained overnight with MMIR1 (5 µg/ml) and MMIR- (20 µg/ml) multiplexed with Hoechst 33342 (0,5 µM) and Sytox green (30 nM). Data shows the average number of viable cells /total cell number ± standard error of 4 replicates. B: Viability assay using CellTiter-Glo Luminescent Cell Viability assay (Promega) shows no statistical cellular toxicity after 17 h incubation of MMIR1 probe (0-50 µg/ml) on live HCT116 cells. Results were normalized by extracting the total cell proteins and BCA assay, showing the average with background subtraction ± standard error of

12 replicates. C: Normalized viability of HCT116 cells, stained with MMIR1 and treated with mitochondrial uncoupler FCCP (4 μM) and inhibitors oligomycin A (10 μM) 5 min before the cell lysis. Data shows the average with background subtraction ± standard error of 4 spheroids. D: Normalized effect of MMIR1 on overall viability (total cell ATP) of cancer and non-cancer cell lines in 2D. Data shows the average with background subtraction ± standard error of 3-5 spheroids E: ATP assay in 3D for HCT116 spheroids, stained with MMIR1 and normalized by their size (area square). Data shows the average with background subtraction ± standard error of 8-12 spheroids.

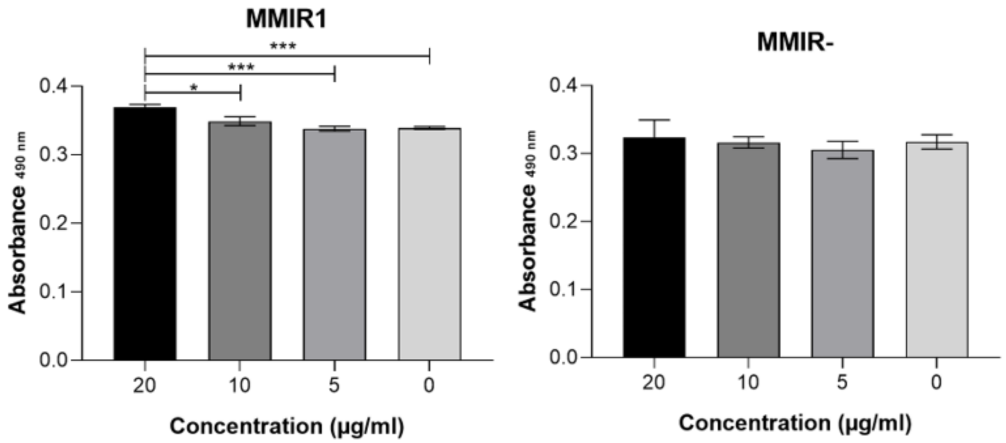


**Supplementary figure S17: Viability assay using the CellTiter 96 Aqueous Non-Radioactive Cell Proliferation Assay (MTS, Promega) shows no cellular toxicity due to 24 h incubation of both cationic (MMIR1) and negatively charged (MMIR-) nanosensors with live HCT116 cells**. Results shown are the average with background subtraction ± standard error of 6 repeats.

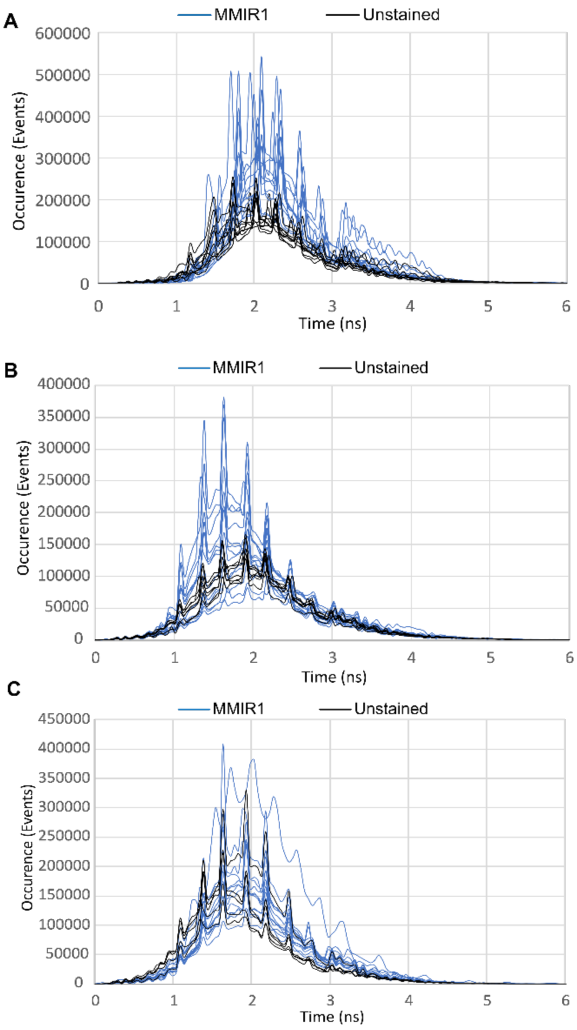


**Supplementary figure S18: Two-photon FLIM of NAD(P)H shows no changes in cell redox (NAD(P)H fluorescence lifetimes) in response to MMIR1 addition to live HCT116 spheroids**. Fluorescence lifetime histograms of unstained (black) and MMIR1 (10 µg/ml, blue)-stained HCT116 spheroids with initial seeding densities of 50 (A), 500 (B), and 10,000 (C) cells per spheroid and 4-5 days growth.

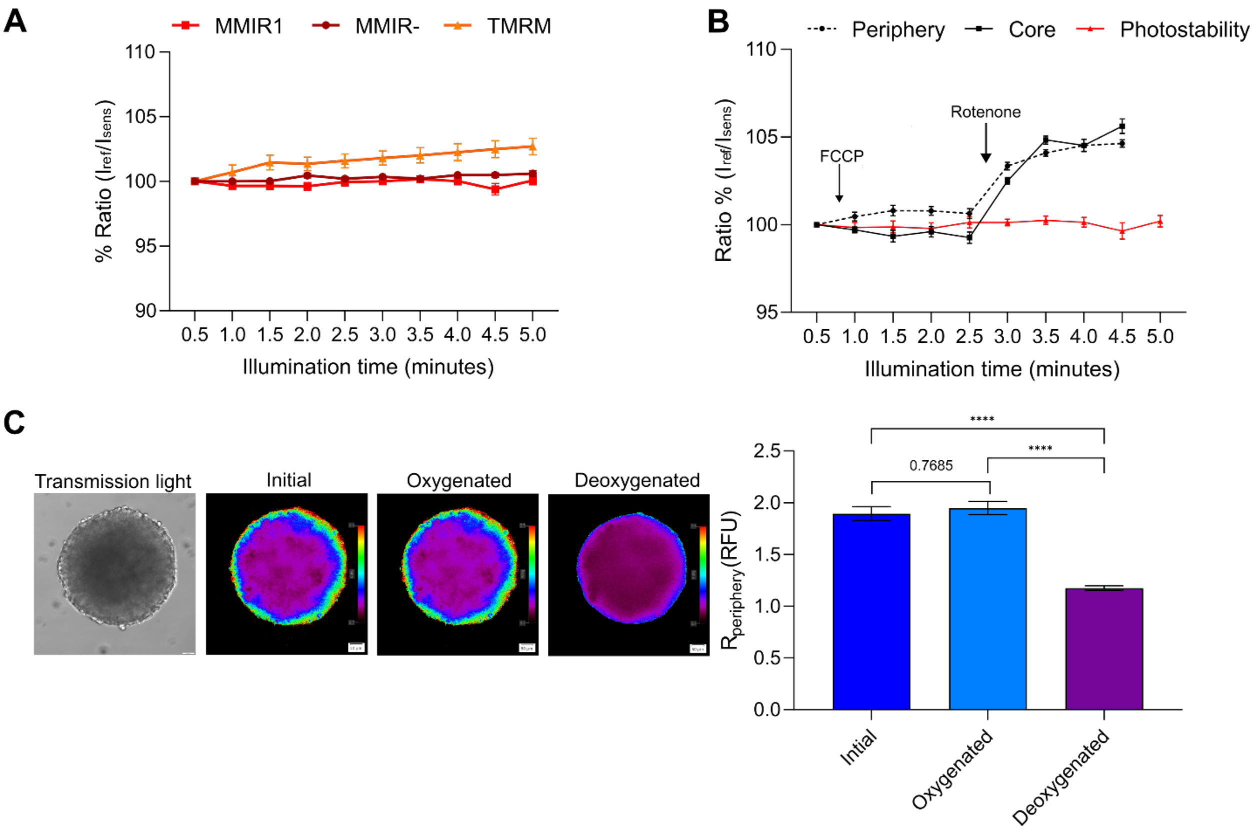


**Supplementary figure S19: Dynamic response of MMIR probes to drugs affecting cell bioenergetics and deoxygenation.** A: Photostability of both MMIR1 and MMIR- in HCT116 spheroids after repeated illumination. Commercially cationic orange-red mitochondrial dye TMRM was used as a reference. N=10 B: Kinetic response of MMIR1 intensity ratio to mitochondrial uncoupler (1 µM FCCP) and mitochondrial inhibitor (1 µM Rotenone) in comparison to photostability kinetics (%). C: Changes in intensity ratio in HCT116 spheroid at oxygenated (1 µM Antimycin A and 1 µM Rotenone) and subsequently deoxygenated (250 ug/ml glucose oxidase and potassium sulfite solution addition). Scale bar 100 µm. Significant difference in intensity of $O_2$-sensitive dye between oxygenated and deoxygenated spheroids. TMRM: Tetramethylrhodamine, methyl ester. ****: $P < 0.0001$.

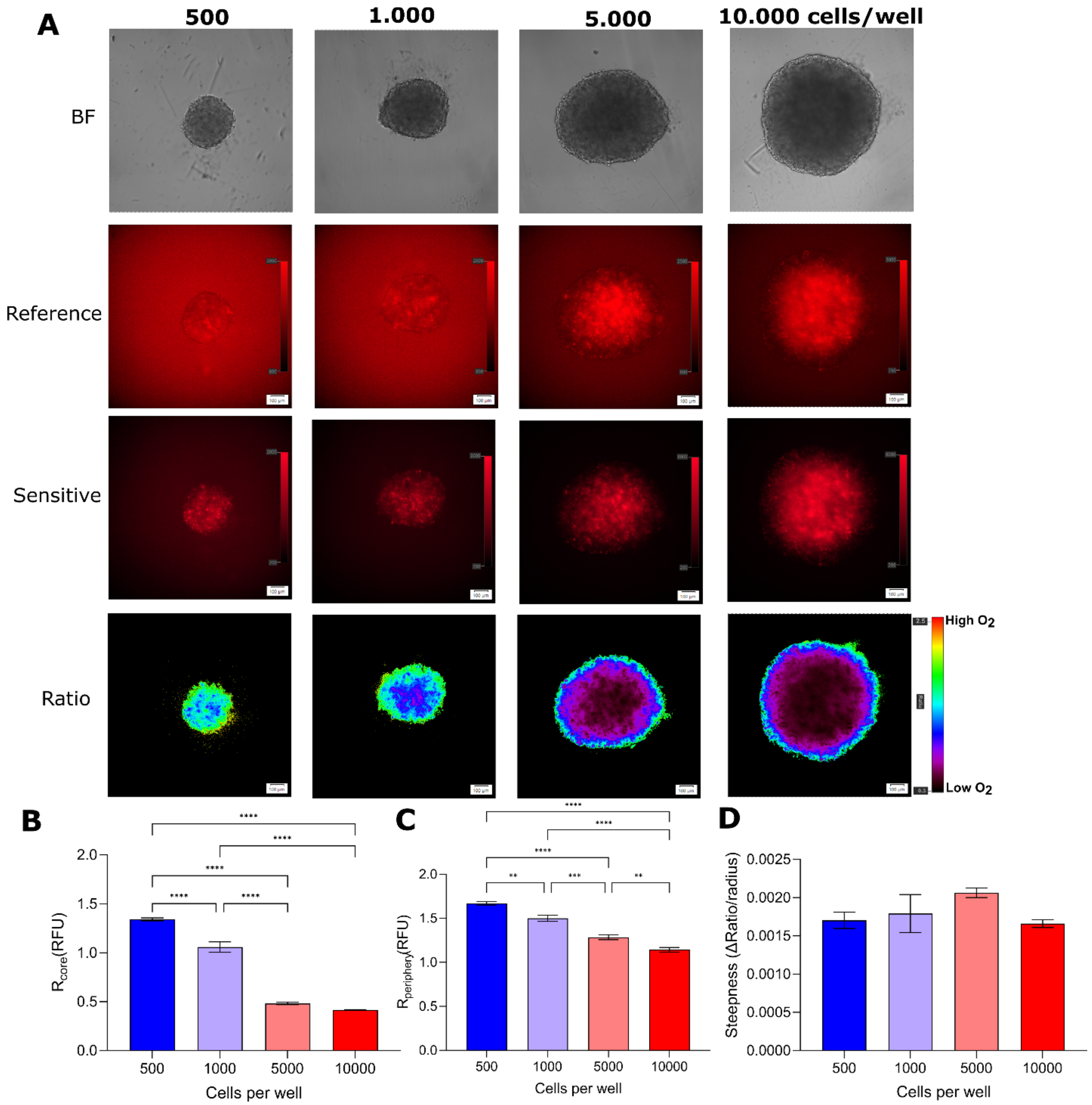


**Supplementary figure S20: Size-dependent oxygenation of live HCT116 cells spheroids.** A: HCT116 cells formed on a lipidure-coated plate in multiple sizes (500, 1,000, 5,000, and 10,000 cells per well) for 5 days, show a decrease in oxygenation in bigger spheroids due to limited oxygenation diffusion. Scale bar is 100 µm. B: MMIR1 ratio measurements at the periphery. C: MMIR1 ratio measurements at the core. D: Steepness of the oxygenation gradient. Results show the average ± standard error of 6 spheroids.

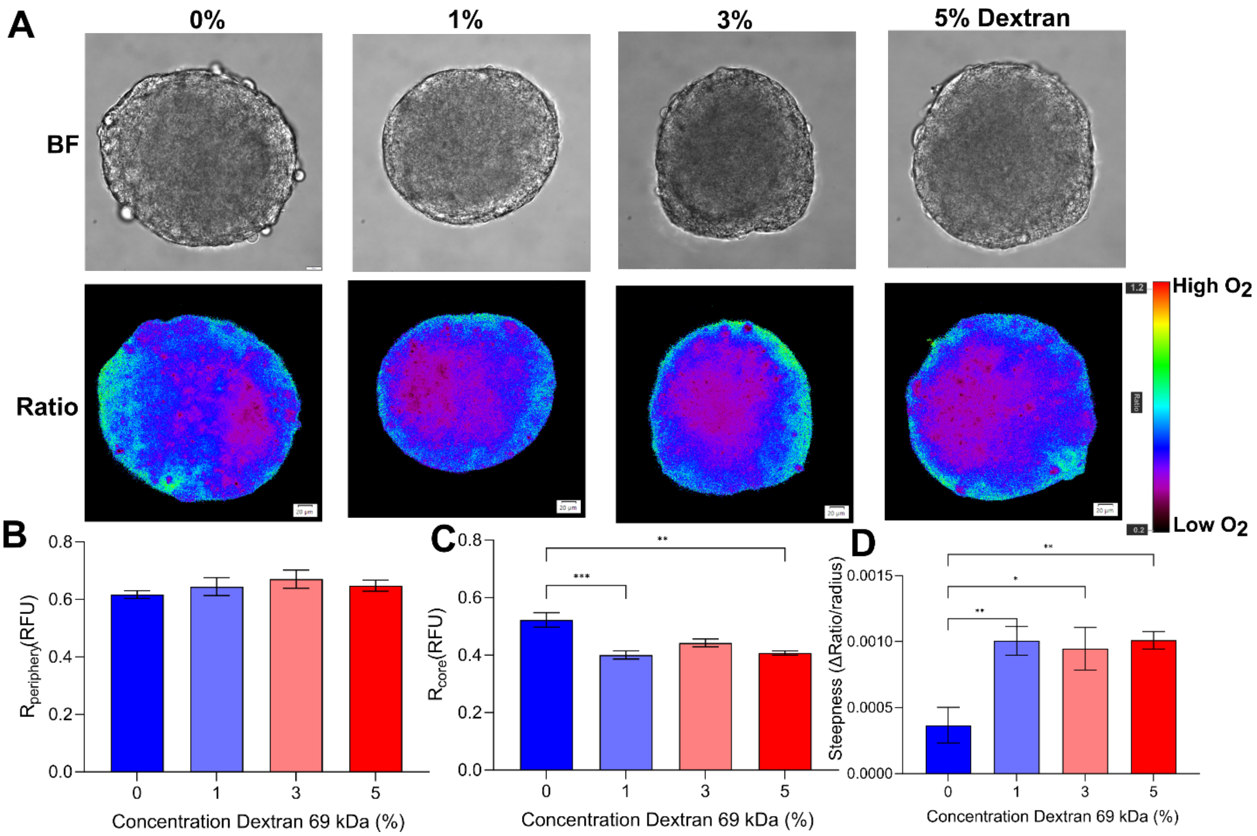


**Supplementary figure S21: Increased cell medium viscosity results in the formation of a more hypoxic core.** A: HCT116 spheroids formed using agarose micromolds were adapted to imaging media containing 0-5% dextran (0.77- 2.25 cP) for 4 h before imaging. Scale bar is 20 µm. B: MMIR1 ratio measurements at the periphery. C: MMIR1 ratio measurements at the core. D: Steepness of the oxygenation gradient. Results show the average ± standard error of 9 spheroids.

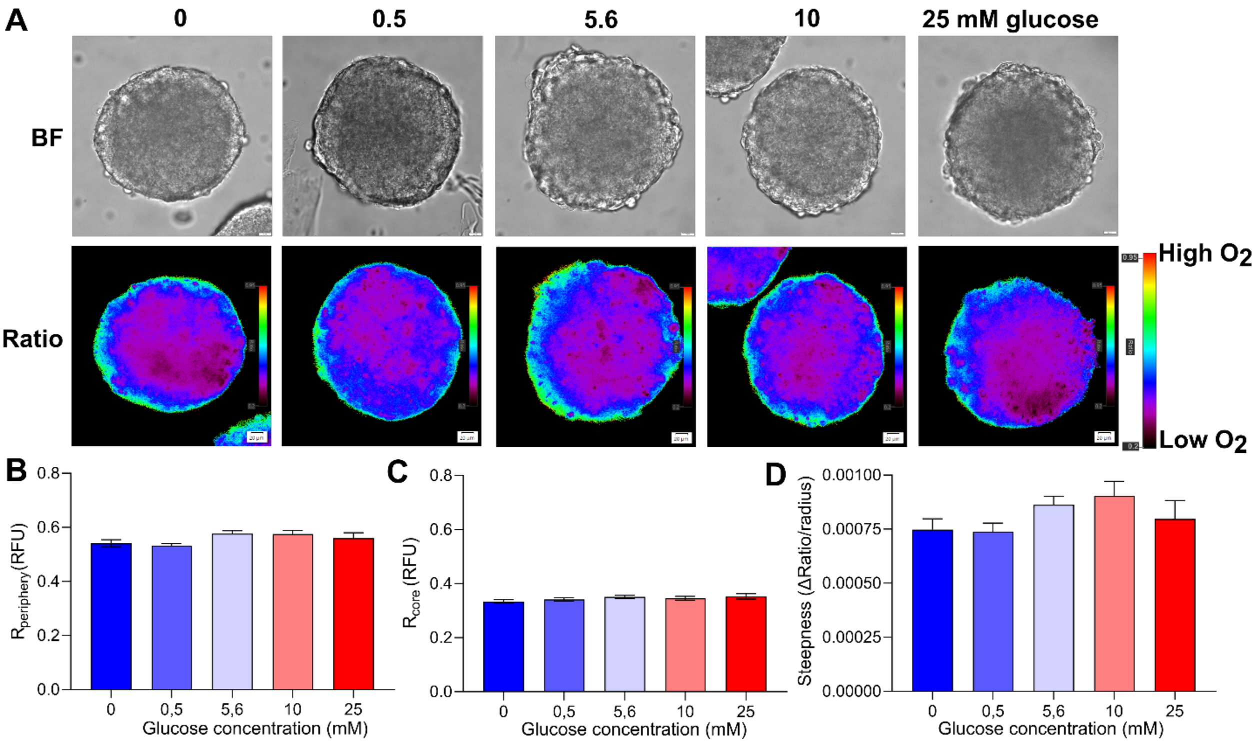


**Supplementary figure S22: Decreased cell media glucose concentration does not affect spheroids oxygenation.** A: HCT 116 spheroids formed using agarose micromolds were adapted to imaging media containing 0-25 mM D(+)-glucose for 4 h before imaging. Scale bar is 20 µm. B: MMIR1 ratio measurements at the periphery. C: MMIR1 ratio measurements at the core. D: Steepness of the oxygenation gradient. Results show the average ± standard error of 9-14 spheroids.

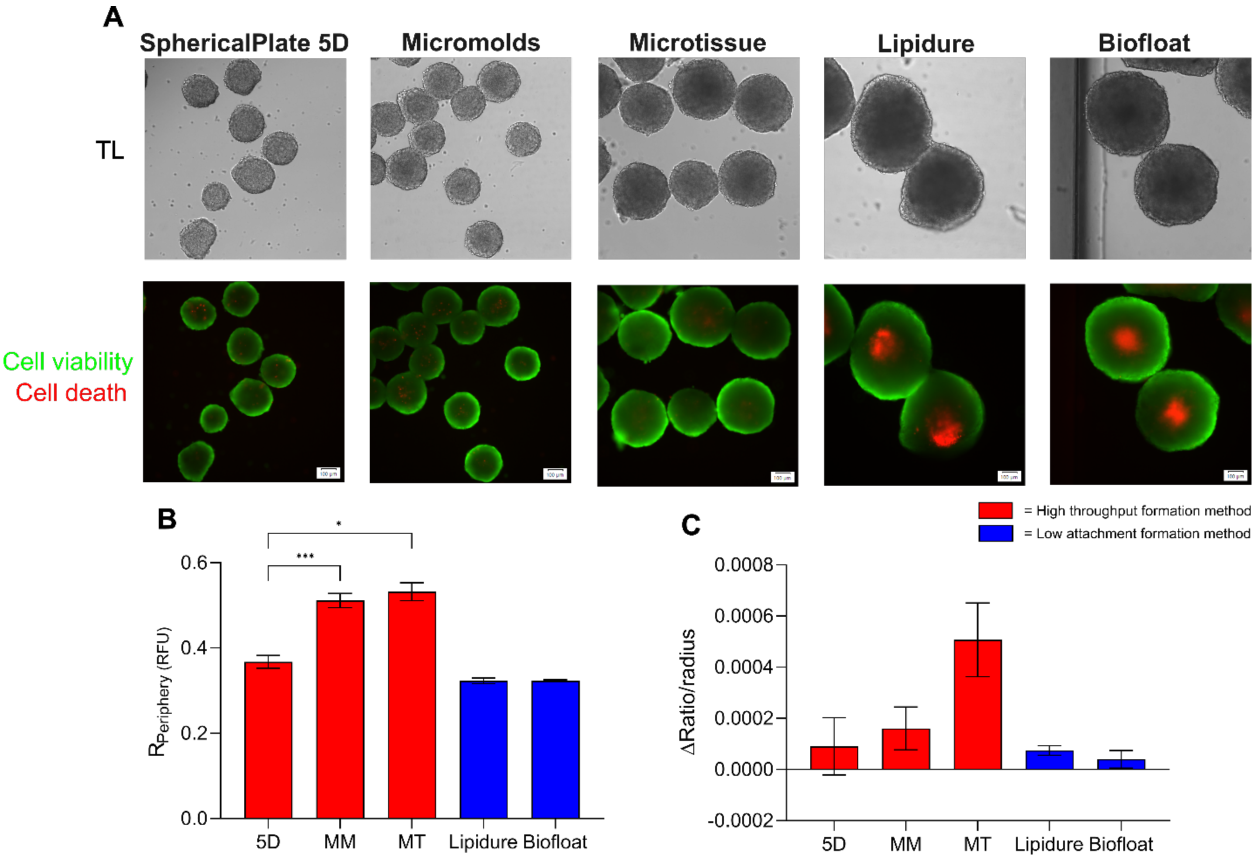


**Supplementary figure S23: Formation methods affect the viability and oxygenation in live HCT116 spheroids.** A: HCT116 spheroids (initial seeding density 500 cells) were grown for 5 days before additional 1 hour-long staining with Propidium Iodide (cell necrosis, red, 1 µg/ml) and Calcein Green (viable cells, green, 1 µg/ml). Low-attachment formation methods lead to bigger spheroids containing a necrotic core (red). Scale bar is 100 µm B: MMIR1 ratio measurements at the periphery show lower oxygenation in 5D SphericalPlate spheroids compared to other high throughput methods, while low attachment methods are similar C: Steepness of the oxygenation gradient is similar in all methods. Red= high throughput formation methods. Blue= low attachment formation methods. Results show the average ± standard error of 6-16 spheroids. 5D= 5D SphericalPlate, MM= Micromold method, MT= MicroTissue method.

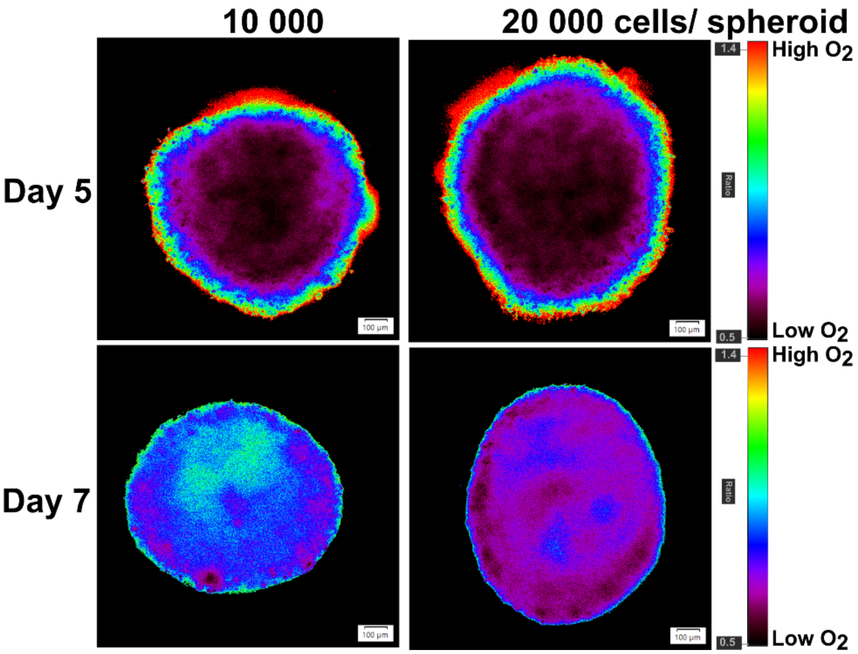


**Supplementary figure S24: Inverted gradient formation in live HCT116 spheroids is both time and size-dependent.** HCT116 spheroids formed by initial seeding 10,000 and 20,000 cells per well are monitored at days 5 and 7. Scale bar is 100 µm.

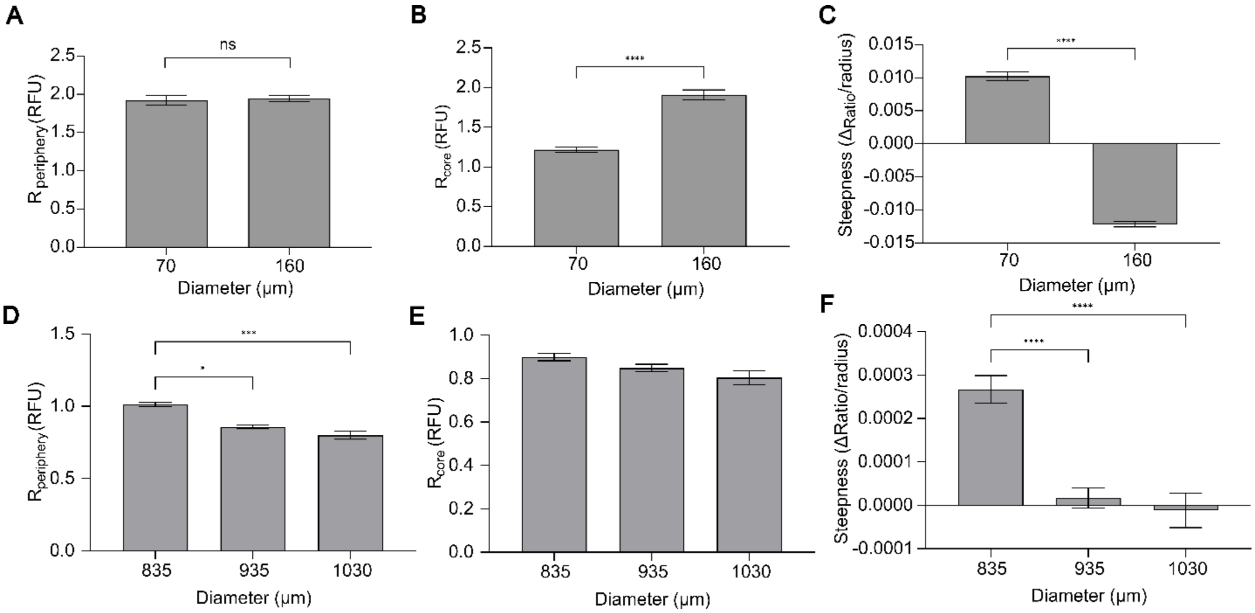


**Supplementary figure S25: Analysis of "inverted" oxygenation gradient in live DPSCs and HCT116 spheroids** (see also **Fig. 5**, main text). A-C DPSCS was formed using microagarose molds and D-F HCT116 was formed using Lipidure®-coated plates. A and D: MMIR1 ratio measurements at the periphery. B and E: MMIR1 ratio measurements at the core. C and F: Oxygenation gradients.

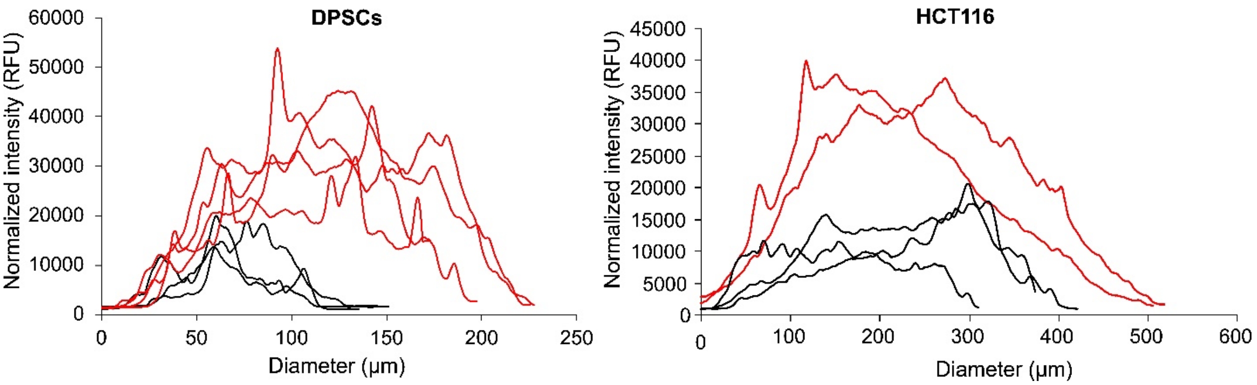


**Supplementary figure S26: Cell death comparison between "direct" and "inverted" gradients in DPSCs and HCT116 spheroids formed on respectively microagarose molds and Lipidure®-coated plates.** Co-staining with propidium iodide, PI (cell death, 1 µg/ml 1h). Data shows line profiles of normalized (by area squared) mean intensity ± SEM of 7 spheroids (DPSCs) and 5 spheroids (HCT116). PI= propidium iodide.

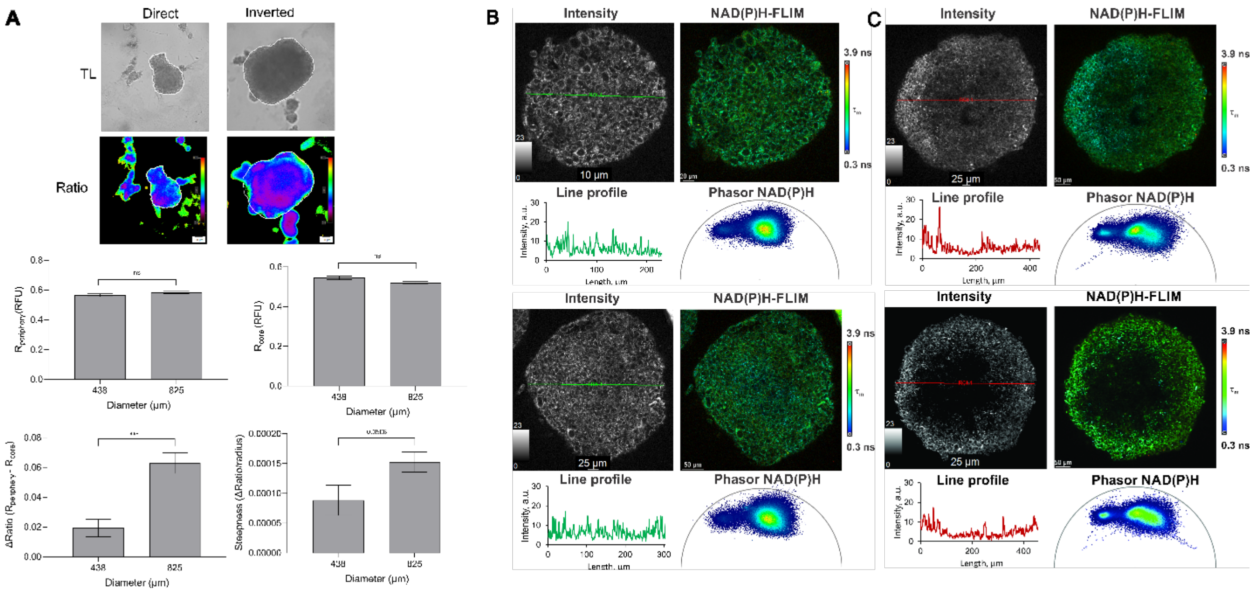


**Supplementary figure S27: Examples of two-photon NAD(P)H-FLIM microscopy and phasor plots of HCT116 spheroids with different sizes. MMIR1 ratiometric analysis, NAD(P)H intensity line profiles and phasor FLIM plots are shown**. A: Ratiometric analysis of MMIR1 probe. B: Direct gradient. C: Indirect gradient.

**Supplementary table ST1:** STR authentication HCT116 cell lines

| Core STR markers | ATCC HCT 116(CCL-247) | HCT 116 HWANG | HCT 116 ODW |
|---|---|---|---|
| D7S820 | 11, 12 | 11,12,13 | 11,12,13 |
| CSF1PO | 7, 10 | 7, 10 | 7,10,11 |
| TH01 | 8, 9 | 8, 9 | 8, 9 |
| D13S317 | 10, 12 | 10,11,12,13 | 10,11,12,13 |
| D16S539 | 11, 13 | 11,12,13,14 | 11,12,13,14 |
| vWA | 17, 22 | 16,17,18,21,22,23 | 16,17,18,21,22,23 |
| TPOX | 8, 9 | 8, 8 | 8, 9 |
| AMEL | X,Y | X,X | X,Y |
| D5S818 | 10, 11 | 10,11,12 | 10,11,12 |

**Supplementary table ST2:** Composition of used growth and imaging media.

| Cell line | Basal medium | Supplements |
|---|---|---|
| DPSC | MEM alpha (Gibco, 32561) | 10% Heat-inactivated FBS (Gibco, 12662) |
| HUVEC | Endothelial Cell Basal Medium 2 (PromoCell, C-22211) | 1x Endothelial Cell Growth Medium 2 SupplementMix (PromoCell, C-39216) |
| HCT116 WT | McCoy's 5A media (VWR, 392-0420) | 10% Heat-inactivated FBS (Gibco, 12662)<br>1 mM Sodium Pyruvate (Gibco, 11360)<br>10 mM HEPES (Gibco, 15630) |
| HCT116 $SCO_2$ $^{-/-}$ | McCoy's 5A media (VWR, 392-0420) | 10% Heat-inactivated FBS (Gibco, 12662)<br>1 mM Sodium Pyruvate (Gibco, 11360)<br>10 mM HEPES (Gibco, 15630) |
| HCT116 KO | McCoy's 5A media (VWR, 392-0420) | 10% Heat-inactivated FBS (Gibco, 12662)<br>1 mM Sodium Pyruvate (Gibco, 11360)<br>10 mM HEPES (Gibco, 15630) |
| SKOV3 | McCoy's 5A media (VWR, 392-0420) | 10% Heat-inactivated FBS (Gibco, 12662)<br>1 mM Sodium Pyruvate (Gibco, 11360)<br>10 mM HEPES (Gibco, 15630) |
| MCF7 | DMEM (Sigma, D5030) | 2 mM GlutaMAX (Gibco, 35050)<br>10 mM HEPES (Gibco, 15630)<br>1 mM Sodium pyruvate (Gibco, 11360)<br>10% Heat-inactivated FBS (Gibco, 12662) |
| PANC1 | DMEM (Sigma, D5030) | 2 mM L-glutamine (Gibco, 25030)<br>10 mM HEPES (Gibco, 15630)<br>1 mM Sodium pyruvate (Gibco, 11360)<br>10% Heat-inactivated FBS (Gibco, 12662) |
| HEK293T | DMEM (Sigma, D5030) | 2 mM L-glutamine (Gibco, 25030)<br>10 mM HEPES (Gibco, 15630)<br>1 mM Sodium pyruvate (Gibco, 11360)<br>10% Heat-inactivated FBS (Gibco, 12662) |
| Imaging medium | DMEM (Sigma, D5030) | 2 mM L-glutamine (Gibco, 25030)<br>10 mM HEPES, pH 7.2 (Gibco, 15630)<br>1 mM Sodium pyruvate (Gibco, 11360)<br>10% Heat-inactivated FBS (Gibco, 12662)<br>10 mM D(+)-glucose (Merck, 8342) |

**Supplementary table ST3:** morphological characterization of spheroid populations

| Number | General $O_2$ gradient shape | Glycolytic core size (% from total spheroid area) | *d*, pixels | Morphological characteristics of individual spheroids |
|---|---|---|---|---|
| 1 | Forward | 0 | 436.75 | Diameter is 275.85 µm. No clear core; homogeneous distribution of long and short autofluorescence lifetime |
| 2 | Forward | 29.84078069 | 400.94 | Diameter is 388.23 µm. Small core with close to central localization |
| 3 | Forward | 22.66822022 | 396.24 | Diameter is 320.81 µm. Small core with close to central localization |
| 4 | Forward | 0 | 407.91 | Diameter is 239.18 µm. No clear core; homogeneous distribution of long and short autofluorescence lifetime |
| 5 | Forward | 25.34300697 | 428.16 | Diameter is 275.97 µm. Small core with close to central localization |
| 6 | Forward | 12.02982783 | 433.12 | Diameter is 348.85 µm. Small core with close to central localization |
| 7 | Forward | 0 | 414.62 | Diameter is 337.24 µm. No clear core; homogeneous distribution of long and short autofluorescence lifetime |
| 8 | Forward | 0 | 451.03 | Diameter is 328.89 µm. No clear core; homogeneous distribution of long and short autofluorescence lifetime |
| **Summary:** | *50 % of spheroids with an average glycolytic core size of 22.5% from the total spheroid area* | | | |
| 1 | Inverted | 26.96279845 | 415.21 | Diameter is 455.18 µm. Core with close to central localization |
| 2 | Inverted | 47.93264855 | 402.10 | Diameter is 415.09 µm. Core shifted toward spheroid periphery |
| 3 | Inverted | 66.73045697 | 377.75 | Diameter 461.6 µm. Core shifted toward spheroid periphery; cells with long lifetime and low intensity observed inside the core |
| 4 | Inverted | 41.66305525 | 402.57 | Diameter is 449.18 µm. Core shifted toward spheroid periphery; cells with long lifetime and low intensity observed inside the core |
| 5 | Inverted | 52.06284481 | 391.30 | Diameter is 451.96 µm. Core shifted toward spheroid periphery; cells with long lifetime and low intensity observed inside the core |
| 6 | Inverted | 71.83703873 | 377.59 | Diameter is 507.7 µm. Core with close to central localization; cells with long lifetime and low intensity observed inside the core, appearance of empty spaces |
| 7 | Inverted | 70.54834055 | 390.93 | Diameter is 615 µm. Core with close to central localization; the drop of autofluorescence intensity in the middle of the core |
| 8 | Inverted | 44.84668557 | 413.01 | Diameter is 564.62 µm. Core with close to central localization; cells with long lifetime and low intensity observed inside the core, appearance of empty spaces |

| | | | | |
|---|---|---|---|---|
| 9 | Inverted | 71.78658835 | 421.19 | Diameter is 577.51 µm. Core shifted toward spheroid periphery; the drop of autofluorescence intensity inside the core |
| 10 | Inverted | 47.52877651 | 405.58 | Diameter is 579.48 µm. Core with close to central localization; cells with long lifetime and low intensity observed inside the core, appearance of empty spaces |
| **Summary:** | *100% spheroids with an average glycolytic core size of 54.2% from the total spheroid area* | | | |
